# Immune Correlates of Protection by mRNA-1273 Immunization against SARS-CoV-2 Infection in Nonhuman Primates

**DOI:** 10.1101/2021.04.20.440647

**Authors:** Kizzmekia S. Corbett, Martha C. Nason, Britta Flach, Matthew Gagne, Sarah O’ Connell, Timothy S. Johnston, Shruti N. Shah, Venkata Viswanadh Edara, Katharine Floyd, Lilin Lai, Charlene McDanal, Joseph R. Francica, Barbara Flynn, Kai Wu, Angela Choi, Matthew Koch, Olubukola M. Abiona, Anne P. Werner, Gabriela S. Alvarado, Shayne F. Andrew, Mitzi M. Donaldson, Jonathan Fintzi, Dillon R. Flebbe, Evan Lamb, Amy T. Noe, Saule T. Nurmukhambetova, Samantha J. Provost, Anthony Cook, Alan Dodson, Andrew Faudree, Jack Greenhouse, Swagata Kar, Laurent Pessaint, Maciel Porto, Katelyn Steingrebe, Daniel Valentin, Serge Zouantcha, Kevin W. Bock, Mahnaz Minai, Bianca M. Nagata, Juan I. Moliva, Renee van de Wetering, Seyhan Boyoglu-Barnum, Kwanyee Leung, Wei Shi, Eun Sung Yang, Yi Zhang, John-Paul M. Todd, Lingshu Wang, Hanne Andersen, Kathryn E. Foulds, Darin K. Edwards, John R. Mascola, Ian N. Moore, Mark G. Lewis, Andrea Carfi, David Montefiori, Mehul S. Suthar, Adrian McDermott, Nancy J. Sullivan, Mario Roederer, Daniel C. Douek, Barney S. Graham, Robert A. Seder

## Abstract

Immune correlates of protection can be used as surrogate endpoints for vaccine efficacy. The nonhuman primate (NHP) model of SARS-CoV-2 infection replicates key features of human infection and may be used to define immune correlates of protection following vaccination. Here, NHP received either no vaccine or doses ranging from 0.3 – 100 μg of mRNA-1273, a mRNA vaccine encoding the prefusion-stabilized SARS-CoV-2 spike (S-2P) protein encapsulated in a lipid nanoparticle. mRNA-1273 vaccination elicited robust circulating and mucosal antibody responses in a dose-dependent manner. Viral replication was significantly reduced in bronchoalveolar lavages and nasal swabs following SARS-CoV-2 challenge in vaccinated animals and was most strongly correlated with levels of anti-S antibody binding and neutralizing activity. Consistent with antibodies being a correlate of protection, passive transfer of vaccine-induced IgG to naïve hamsters was sufficient to mediate protection. Taken together, these data show that mRNA-1273 vaccine-induced humoral immune responses are a mechanistic correlate of protection against SARS-CoV-2 infection in NHP.

**One-Sentence Summary:** mRNA-1273 vaccine-induced antibody responses are a mechanistic correlate of protection against SARS-CoV-2 infection in NHP.

---

Severe acute respiratory syndrome coronavirus 2 (SARS-CoV-2), the causative agent of coronavirus disease 2019 (COVID-19), has led to more than 138 million infections and 3 million deaths worldwide as of April 15, 2021 (*1*). Mass vaccination offers the most efficient public health intervention to control the pandemic. Two mRNA-based vaccines, Moderna’s mRNA-1273 and Pfizer/BioNTech’s BNT 162b2, both of which encode the prefusion-stabilized spike glycoprotein (S-2P) (*2, 3*), showed >94% efficacy against symptomatic COVID-19 in interim Phase 3 analyses (*4, 5*) and are currently being administered globally. Several other vaccines have shown 60-80% efficacy against COVID-19 in Phase 3 trials (*6, 7*), and a number of candidate vaccines are in earlier stages of clinical development (*8*). A critical issue for optimizing the use of COVID-19 vaccines is defining an immune correlate of protection. This surrogate of vaccine efficacy can be used to inform potential dose reduction, advance approval of other vaccine candidates in lieu of Phase 3 efficacy data, extend indications for use to other age groups, and provide insights into the immune mechanisms of protection (*9*).

The nonhuman primate (NHP) model has been used to demonstrate immunogenicity and protective efficacy against SARS-CoV-2 with a number of vaccine candidates (*10–13*). The high level of protection achieved with mRNA vaccines in NHP using clinically relevant dose regimens has been consistent with results from human trials. This model exhibits upper and lower airway infection and pathology similar to clinical presentations of mild COVID-19 disease in humans (*14*). While assessment of immune correlates of viral load after primary infection has been completed in NHP (*15*), there are no studies to date that have specifically defined immune correlates of protection in upper and lower airways after vaccination with any product approved for use in humans.

We used immunogenicity and protection assessments from our previous NHP mRNA-1273 vaccine study (*13*) to hypothesize that serum antibody measurements serve as immune correlates of protection. Here, in a dose de-escalation study, we evaluated how multiple measurements of humoral and cellular immunity correlate with the reduction of viral replication in the upper and lower airway following challenge. Antibody analyses were also performed on bronchoalveolar lavages (BAL) and nasal washes after vaccination to assess site-specific immune correlates. Last, we demonstrated the ability of passively transferred IgG from mRNA-immunized NHP to mediate protection in a highly pathogenic Syrian hamster SARS-CoV-2 challenge model. Together, these studies support spike (S)-specific antibodies as a correlate of protection, highlight the ability of localized mucosal antibodies to control upper and lower airway viral replication, and confirm mRNA-1273-induced IgG to be sufficient for protection against SARS-CoV-2 infection in preclinical models.

## Results

### mRNA-1273 vaccination elicits robust antibody responses in a dose-dependent manner

We previously demonstrated dose-dependency of serum antibody responses in NHP following vaccination with 10 or 100 μg of mRNA-1273, with high-level protection against SARS-CoV-2 challenge in both dose groups (Fig. S1A) (*13*). These and other immunogenicity outcomes from an additional NHP study in which animals were vaccinated with 30 μg of mRNA-1273 (Fig. S1B) were used to design a study to evaluate immune correlates of protection following mRNA-1273 vaccination in the current study (Fig. S1C). Doses of mRNA-1273 ranging from 0.3 to 30 μg were administered in the standard primary regimen at weeks 0 and 4 to generate a range of immune responses and protective outcomes.

We first assessed temporal serum S-specific antibody binding or avidity and pseudovirus neutralization responses post-prime and -boost. Consistent with our previous report (*13*), S-specific IgG binding to the conformationally defined prefusion S-2P protein (*2, 3*) was increased over baseline after each immunization, reaching 7,900 and 64,000 median reciprocal endpoint titers by 4 weeks post-prime and -boost, respectively, following immunization with 30 μg of mRNA-1273 (Fig. S2A). There was an 8-10-fold increase in S-specific binding antibodies after the boost in all dose groups, except for the 0.3 μg dose for which boosting elicited 300-fold more S-specific antibodies. S-specific antibody avidity was also increased, by 2-fold, after the boost in all vaccine groups except for the 0.3 μg dose, and there were no differences between the vaccine groups (Fig. S2B). D614G pseudovirus neutralizing antibody responses 4 weeks post-prime were only detectable in the 30 μg dose group (median reciprocal ID_50_ = 76), increasing by ~1 log_10_ post-boost (Fig. S2C).

For analyses of immune correlates, we used data from six different qualified antibody assays performed at the time of SARS-CoV-2 challenge, 4 weeks post-boost. Anti-S-specific (Fig. 1A) and anti-receptor binding domain (RBD) (Fig. 1B) responses were measured using Meso-Scale Discovery Multiplex ELISA, validated for use in Phase 3 clinical SARS-CoV-2 vaccine trials; here, antibody binding titers are reported in international units (IU) defined by World Health Organization (WHO) standards. Binding antibody titers increased compared to control animals in a dose-dependent manner ranging from a median of 55 to 5,800 IU/mL at 0.3 and 100 μg, respectively, for S-specific IgG, and 66 to 10,400 IU/mL for RBD-specific IgG (Fig. 1A-B). There was also a dose-dependent reduction in median ACE2 binding inhibition comparing 100 μg to 1 μg of mRNA-1273 (Fig. 1C), reaching a maximum of 270-fold. *In vitro* neutralizing activity was determined using three orthogonal assays. First, for the lentiviral-based D614G pseudovirus neutralization assay qualified for use in Phase 3 clinical studies, there was a dose-dependent decrease with a median reciprocal ID_50_ titer of 23,000 at the 100 μg dose and 49 following immunization with 1 μg of mRNA-1273 (Fig. 1D). VSV-based pseudovirus (Fig. 1E) and live virus (Fig. 1F) neutralization followed the same significant dose-dependency trend. Assessments of antibody binding and neutralization responses were highly correlated with one another, suggesting mRNA-1273 immunization elicits high titer S-binding antibody responses and high-level functional antibody responses (Fig. 2).

**Fig. 1.**
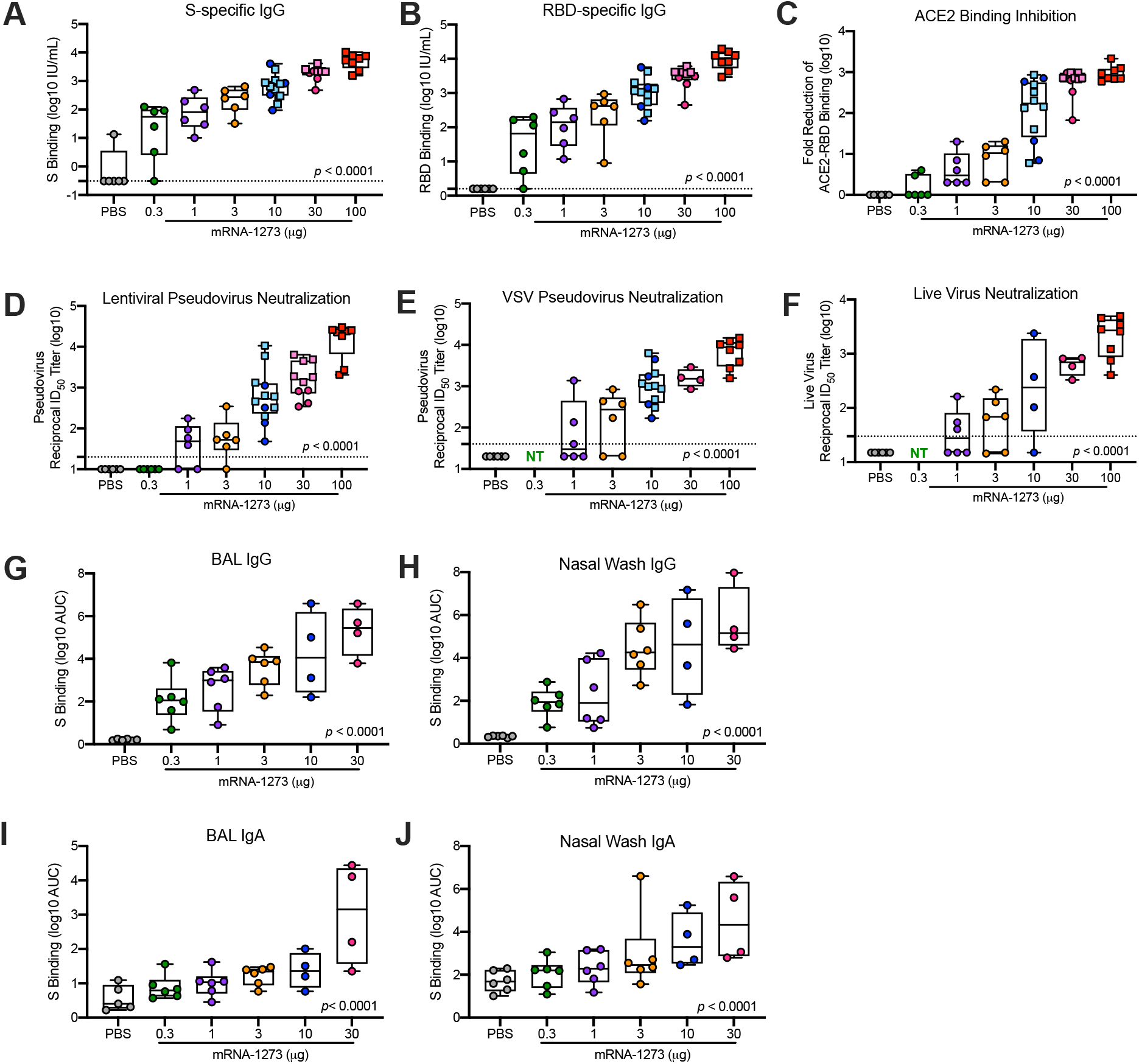
Antibody responses following mRNA-1273 immunization. Rhesus macaques were immunized according to Fig. S1 with PBS (gray) or mRNA-1273 (0.3 μg – green, 1 μg – purple, 3 μg – orange, 10 μg – blue, 30 μg – pink, or 100 μg – red). Sera collected 4 weeks post-boost, immediately before challenge, were assessed for SARS-CoV-2 S-specific (A) and RBD-specific (B) IgG by MULTI-ARRAY ELISA, inhibition of ACE2 binding to RBD (C), SARS-CoV-2 lentiviral-based pseudovirus neutralization (D), SARS-CoV-2 VSV-based pseudovirus neutralization (E), and SARS-CoV-2 EHC-83E focus reduction neutralization (F). BAL (G, I) and nasal washes (H, J) collected 2 weeks post-boost were assessed for SARS-CoV-2 S-specific IgG (A-B) and IgA (C-D) by MULTI-ARRAY ELISA. Squares represent NHP in previous experiments (S1A, VRC-20-857.1 and S1B, VRC-20-857.2); circles represent individual NHP in experiment S1C, VRC-20-857.4. Boxes and horizontal bars denote the IQR and medians, respectively; whisker end points are equal to the maximum and minimum values. Dotted lines indicate assay limits of detection, where applicable. NT = not tested. All measures were significantly correlated with dose (*p*<0.0001), as determined by a test of Spearman’s correlation.

**Fig. 2.**
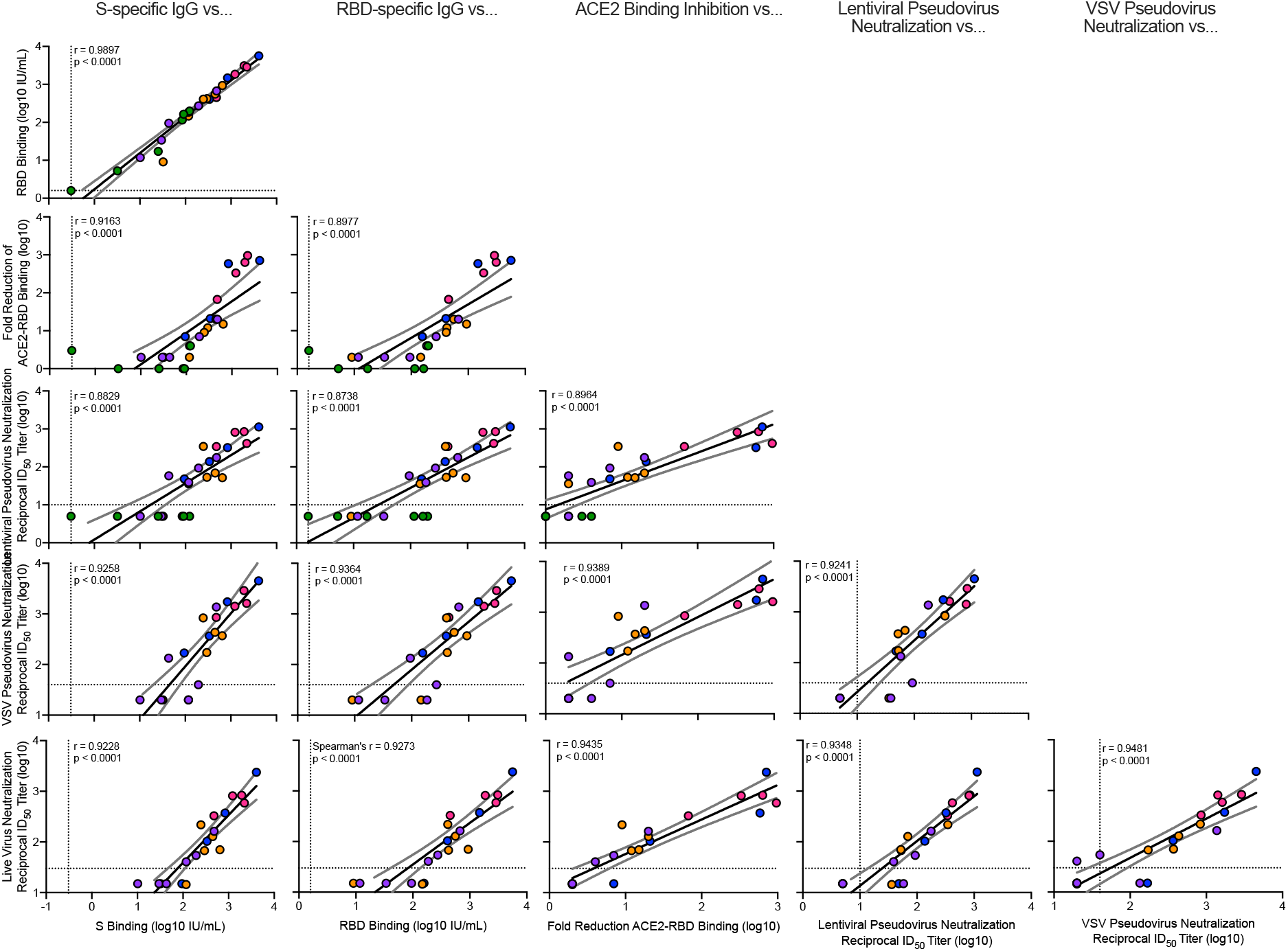
Correlations of humoral antibody analyses. Rhesus macaques were immunized according to Fig. S1C. Plots show correlations between SARS-CoV-2 S-specific IgG, RBD-specific IgG, ACE2 binding inhibition, lentiviral-based pseudovirus neutralization, VSV-based pseudovirus neutralization, and EHC-83E focus reduction neutralization at 4 weeks post-boost. Circles represent individua l NHP, where colors indicate mRNA-1273 dose as defined in Fig. S1C. Dotted lines indicate assay limits of detection. Black and gray lines indicate linear regression and 95% confidence interval, respectively. ‘r’ represents Spearman’s correlation coefficients, and ‘p’ the corresponding p-values.

Given the increasing circulation of SAR-CoV-2 variants of concern, some of which have shown a significant reduction in neutralization sensitivity to vaccine-elicited and convalescent sera (*16–20*), we also assessed the ability of mRNA-1273 immune NHP sera to neutralize two of the SAR-CoV-2 variants of concern. Live viral neutralization of the B.1.1.7 variant (*21*), which is highly transmissible and currently circulating around the world (*22*), was not appreciably decreased as compared to D614G (Fig. S3A). For the B.1.351 variant, which contains multiple mutations in RBD and NTD and shows the greatest reduction of neutralization by vaccine sera (*16, 23, 24*), there was a 9-fold reduction compared to D614G in the 100 μg mRNA-1273 dose group. Notably, 9 of 12 animals immunized with 30 or 100 μg of mRNA-1273 had reciprocal ID_50_ titers > 100, while only 1 of 4 animals in the 10 μg dose group had detectable neutralization activity to B.1.351 (Fig. S3B). The reduction in B.1.351 neutralization capacity of mRNA-1273-induced antibodies mirrors what has been previously shown in NHP or humans using only 30 or 100 μg (*16, 20*), but these data further suggest that mRNA-1273 dose may have a profound effect on eliciting neutralizing antibodies against the B.1.351 variant.

### mRNA-1273 vaccination elicits upper and lower airway antibodies

To provide additional immune data on correlates of protection at the site of infection, antibody responses in the lower and upper airway were assessed from BAL and nasal wash samples, respectively, at 2 weeks post-boost. There was a dose-dependent increase in BAL and nasal wash S-specific IgG and IgA following two doses of mRNA-1273 (Fig. 1G-J). BAL S-specific IgG titers following 0.3 and 30 μg of mRNA-1273 ranged from a median of 110 to 280,000 area under the curve (AUC) (Fig. 1G), and nasal wash S-specific IgG titers ranged from 86 to 142,200 AUC (Fig. 1H). For S-specific IgA, the dose-dependent trend was similar albeit to lower titers where 30 μg of mRNA-1273 elicited 1400 and 21,300 AUC IgA in BAL (Fig. 1I) and nasal washes (Fig. 1J), respectively. Additionally, upper and lower airway antibody responses correlated with each other and the two Phase 3 qualified humoral antibody measurements, S-specific IgG and lentiviral-based pseudovirus neutralization activity; the one exception was that there was no correlation with BAL and nasal wash S-specific IgA (Fig. S4). In all, mRNA-1273 vaccination elicits S-specific IgG and IgA antibodies in both the upper and lower airways, which potentially provide immediate protection at the site of infection.

### mRNA-1273 vaccination elicits S-specific CD4 T cell responses

S-specific CD4 and CD8 T cell responses were assessed 2 weeks post-boost. A direct correlation between dose and Th1 responses was observed (*p*=0.006), where all animals in the 30 μg dose group had Th1 responses (Fig. S5A). In contrast, Th2 responses were low to undetectable in all vaccine dose groups (Fig. S5B). CD8 T cells at these doses of mRNA-1273 were also undetectable. Given the importance of T follicular helper (Tfh) in regulating antibody responses, we extended the analysis to S-specific Tfh cells that express the surface marker CD40L or the canonical cytokine IL-21. Most vaccinated animals had S-specific CD40L+ CD4 Tfh cell responses - the magnitude of which was directly correlated with dose (*p*<0.001) (Fig. S5C). A direct correlation between dose and magnitude of S-specific IL-21 Tfh cell responses was also observed (*p*=0.010). (Fig. S5D). Consistent with previous results (*13, 25, 26*), these data show that mRNA-1273 induced Th1- and Tfh-skewed CD4 responses.

### mRNA-1273 vaccination protects against upper and lower airway SARS-CoV-2 replication

To evaluate the protective efficacy of mRNA-1273 vaccination, all animals in experiment VRC-20-857.4 (Fig. S1C) were challenged 4 weeks post-boost with a total dose of 8×10^5^ PFU of a highly pathogenic stock of SARS-CoV-2 (USA-WA1/2020) by combined intranasal and intratracheal routes for upper and lower airway infection, respectively. This challenge dose was chosen to induce viral loads similar to or higher than those detected in nasal secretions of humans following SARS-CoV-2 infection (*27*). The primary efficacy endpoint analysis used subgenomic RNA (sgRNA) qRT-PCR for the nucleocapsid (N) gene (Fig. 3). N sgRNA is the most highly expressed sgRNA species as a result of discontinuous transcription and thus provides greater sensitivity than the envelope (E) gene (Fig. S6) (*28*), which is most commonly used in other NHP SARS-CoV-2 vaccine studies (*13*) to quantify replicating virus.

**Fig. 3.**
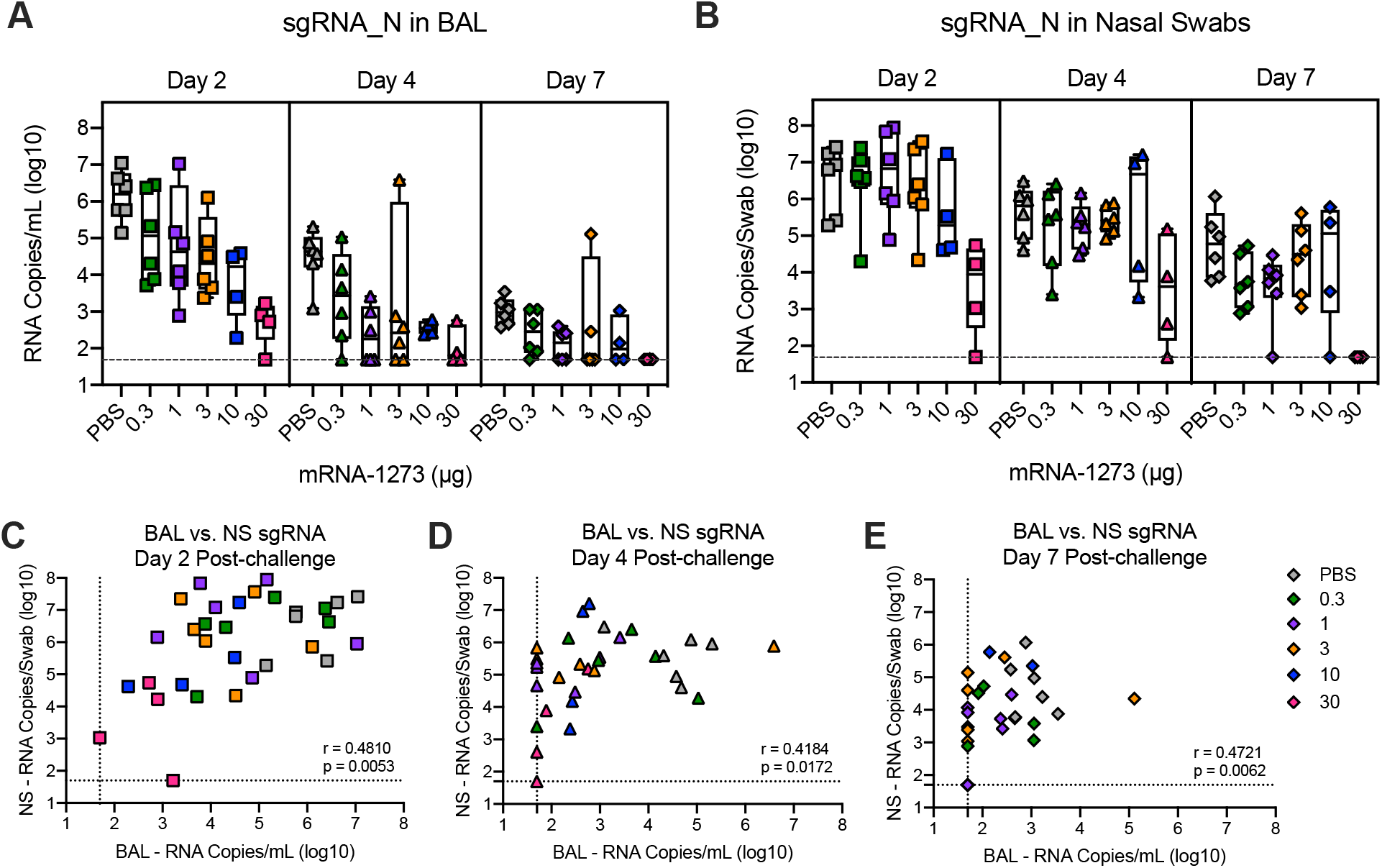
Efficacy of mRNA-1273 against upper and lower respiratory viral replication. Rhesus macaques were immunized and challenged as described in Fig. S1C. BAL (A) and nasal swabs (NS) (B) were collected on days 2 (squares), 4 (triangles), and 7 (diamonds) post-challenge, and viral replication was assessed by detection of SARS-CoV-2 N-specific sgRNA. (A-B) Boxes and horizontal bars denote the IQR and medians, respectively; whisker end points are equal to the maximum and minimum values. (C-E) Correlations shown between BAL and NS sgRNA at days 2 (C), 4 (D), and 7 (E) post-challenge are Spearman’s correlation coefficients (r) and corresponding p-values. Symbols represent individual NHP and may overlap, ie. n=6 animals plotted at assay limit (dotted line) for both BAL and NS in (E).

We observed a vaccine dose effect for protection against viral replication in the upper and lower airway. On days 2 and 4 post challenge, there were ~2 and 5 log_10_ reductions in sgRNA_N in BAL compared to control animals at doses of 1 μg and 30 μg, respectively (Fig. 3A). Moreover, by day 4 post-challenge, the majority of animals vaccinated with 1 μg or higher had low to undetectable sgRNA_E in BAL (Fig. S6A). By contrast, the reduction in sgRNA in nasal swabs was primarily limited to animals receiving 30 μg of mRNA-1273 as compared to control animals (Fig. 3B, Fig. S5B). These data highlight differences in immune responses required for reduction in viral replication for upper and lower airway protection. Post-challenge, there was a strong correlation between sgRNA in the upper and lower airways; however, the virus was more rapidly cleared from the BAL compared to the nasal swab samples. Thus, there was a time-dependent loss of concordance in the correlations with upper and lower airways samples (Fig. 3C-E), suggesting distinct mechanisms for viral clearance in the two compartments.

### mRNA-1273-vaccinated NHP have limited virus and inflammation in lungs

Animals in each of the dose groups were assessed for detection of virus in the lung and histopathology 7- or 8-days post SARS-CoV-2 challenge. In the control animals, SARS-CoV-2 infection caused moderate to severe inflammation that often involved the small airways and the adjacent alveolar interstitium consistent with previous reports (*29–31*). Alveolar air spaces occasionally contained inflammatory cell infiltrates, alveolar capillary septa were moderately thickened, and moderate and diffuse type II pneumocyte hyperplasia was observed. Multiple pneumocytes in the lung sections from the control group were positive for SARS-CoV-2 viral antigen by immunohistochemistry (IHC) (Fig. S7, Table S1). Viral antigen was detected in both control animals but only sporadically across vaccinated animals in various dose groups (Table S1). These observations show that NHP develop mild inflammation in the lung over 1 week following SARS-CoV-2 infection and that vaccination limits or completely prevents inflammation or detection of viral antigen in the lung tissue.

### Post-challenge anamnestic antibody responses are increased in low dose vaccine groups

Following SARS-CoV-2 challenge, we assessed antibody responses in blood, BAL, and nasal washes for up to 28 days to determine if there were anamnestic or primary responses to S or N proteins, respectively (Fig. S8). This analysis provides a functional immune assessment of whether the virus detected in the upper and lower airways by PCR following challenge is sufficient to boost vaccine-induced S-specific antibody responses or elicit primary N responses. In sera, there was no post-challenge increase in S-specific (Fig. S8A), RBD-specific (Fig. S8B), or neutralizing antibodies (Fig. S8C) in the 3, 10, or 30 μg dose groups. In contrast, at doses below 1 μg, there were increased primary S-specific (Fig. S8A), RBD-specific (Fig. S8B), and neutralizing antibody responses (Fig. S8C) at day 28 post-challenge compared to pre-challenge. Similar primary S-specific antibody response trends were also apparent with BAL and nasal wash IgG and IgA responses (Fig. S9). Of note, in comparing pre-challenge N-specific IgG responses to those post-challenge, we only observed seroconversion in the control animals and animals immunized with <3 μg of mRNA-1273 (Fig. S8D).

The reduction of viral replication as determined by sgRNA coupled with limited pathology in the lung and no detectable anamnestic S responses or induction of primary responses to N provide three distinct measures suggesting that vaccine-elicited immune responses, particularly at high doses, were protective. To understand this further, and to establish immune correlates of protective immunity, we explored relationships between immune parameters and viral load.

### Antibody responses correlate with protection against SARS-CoV-2 replication

Prior to conducting study VRC-20-857.4 (Fig. S1C), we pre-specified that our analysis of a potential correlate would focus primarily on the relationship between S-specific binding antibodies and sgRNA levels in NS. Correlations with sgRNA levels in BAL served as an important secondary analysis. The pre-defined primary hypothesis of the study was that S-specific IgG at 4 weeks post-boost (pre-challenge) would inversely correlate with viral replication in the NS at day 2 post-challenge and that vaccine dose may not be predictive of viral replication after adjustment for S-specific IgG. The hypotheses were analogous for the relationship between S-specific IgG at 4 weeks post-boost and day 2 BAL sgRNA.

S-specific IgG at week 8 correlated strongly with sgRNA in both the NS (*p*=0.001) (Fig. 4G, Table S2) and BAL (*p*<0.001) (Fig. 4A, Table S2) at day 2. As shown in Table S2, a 1 log_10_ change in S-specific IgG corresponds to a 1 log_10_ change in sgRNA at day 2 in the NS, and a 0.9 log_10_ change in the sgRNA in the BAL at day 2. Once the S-specific IgG was included in a linear model predicting sgRNA, including dose in the model did not substantially increase the adjusted R^2^, nor was the coefficient significant (*p*=0.115 for NS and *p*=0.214 for BAL). This suggests that the effect of dose on day 2 sgRNA in NS and BAL is fully captured by the adjustment for S-specific IgG and that, in this model, S-specific IgG meets our pre-specified criteria to be considered as a correlate of sgRNA levels in NS and BAL.

**Fig. 4.**
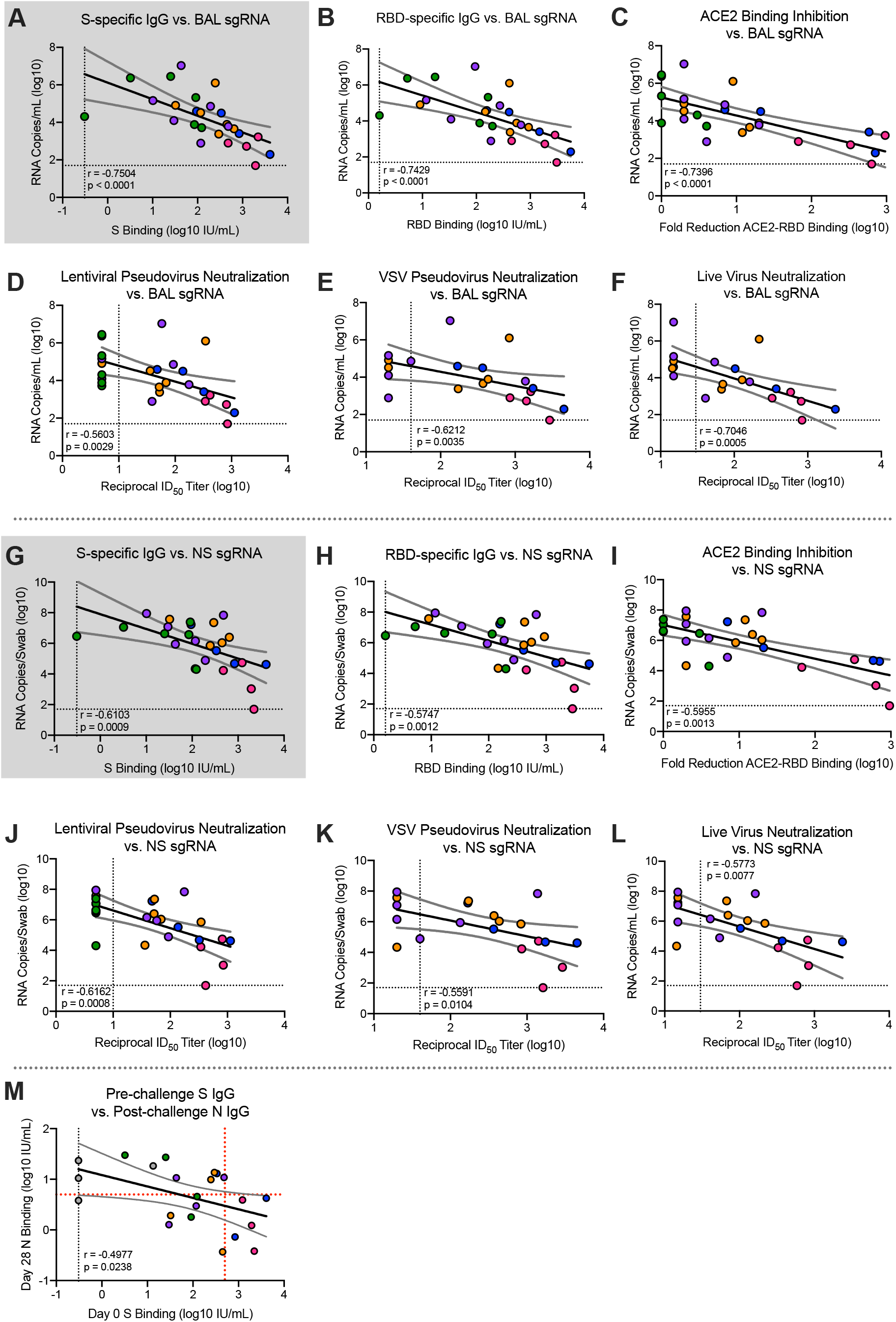
Antibody correlates of protection. Rhesus macaques were immunized and challenged as described in Fig. S1C. Plots show correlations between SARS-CoV-2 N-specific sgRNA in BAL (A-F) and NS (G-L) at day 2 post-challenge and pre-challenge (week 4 post-boost) SARS-CoV-2 S-specific IgG (A, G), RBD-specific IgG (B, H), ACE2 binding inhibition (C, I), SARS-CoV-2 lentiviral-based pseudovirus neutralization (D, J), SARS-CoV-2 VSV-based pseudovirus neutralization (E, K) and SARS-CoV-2 EHC-83E focus reduction neutralization (F, L). Gray shading for S-specific IgG represents the use of this assessment as primary predictor of protection outcome as stated in primary hypothesis. (M) Plot shows correlation between pre-challenge (week 4 post-boost) SARS-CoV-2 S-specific IgG with day 28 post-challenge SARS-CoV-2 N-specific IgG. Circles represent individual NHP, where colors indicate mRNA-1273 dose. Dotted lines indicate assay limits of detection. Black and gray lines indicate linear regression and 95% confidence interval, respectively. In (M), red dotted horizontal line represents 6, the maximum of all pre-challenge values across all groups, and the red dotted vertical line represents a reciprocal S-specific IgG titer of 500, above which none of the animals had day 28 N Binding titers above 6. ‘r’ represents Spearman’s correlation coefficient, and ‘p’ the corresponding p-value.

As RBD-specific IgG, ACE2 binding inhibition, pseudovirus neutralization, and live virus neutralization correlated with S-specific IgG (Fig. 2), analyses of these as potential correlates of sgRNA were also planned. All six antibody measurements were highly correlated with each other (Fig. 2), with vaccine dose (Fig. 1), and with sgRNA in BAL (Fig. 4AF) and NS (Fig. 4G-L). As shown in Table S2A, for all six antibody measurements, dose was not significantly predictive of sgRNA in the BAL after adjusting for antibody levels; for NS, dose remained significantly predictive after adjusting for VSV-based pseudovirus neutralization and marginally significant after adjusting for live virus neutralization. This suggests that in addition to S-specific IgG, RBD-specific IgG, ACE2 binding inhibition, and lentiviral-based pseudovirus neutralization meet our criteria for potential correlates of protection. Furthermore, lower and upper airway S-specific antibodies in the BAL and NS negatively correlated with BAL (Fig. S10A-B) and NS sgRNA levels, respectively (Fig. S10C-D).

To assess the robustness of these findings, these analyses were repeated using logistic regression to model the probability that the sgRNA was below a threshold, defined as 10,000 sgRNA copies for BAL and 100,000 sgRNA copies for NS. These thresholds were chosen to be below all of the sgRNA values in the control animals and within the range of the values for the mRNA-1273-vaccinated animals. The results of these analyses were similar to the primary analyses done on the (log) linear models. In these data, no animal with S-specific IgG >336 IU/mL had BAL sgRNA >10,000 copies/mL (Fig. 4A), and no animal with S-specific IgG >645 IU/mL had NS sgRNA >100,000 copies/swab (Fig. 4G). Last, no animals with a S-binding binding titer of >488 IU/mL had higher N-specific primary antibody responses post-challenge above the background value at the time of challenge; consistent with that, there was a strong negative correlation between pre-challenge S-specific antibodies and post-challenge N-specific antibodies (Fig. 4M). Additionally, there was limited to no lung pathology or viral antigen detection in animals with <10,000 sgRNA copies/mL in BAL, providing additional evidence that mRNA-1273-vaccinated animals were protected from lower airway disease.

We also examined the correlations between T cell responses and sgRNA and found that CD40L+ Tfh cells and any Th1 response were each univariately associated with reduced sgRNA in both BAL and NS. After adjustment for S-specific IgG, none of these remained significantly associated with sgRNA levels in the BAL, suggesting that these T cell measures do not predict sgRNA independently of the binding antibody measured in BAL. However, IL-21+ Tfh, CD40L+ Tfh, and any Th1 response remained significantly predictive of sgRNA levels in NS (Table S2B) further confirming that clearance of virus from BAL and NS have distinct immunological requirements (Fig. 3C-E).

### Passively transferred mRNA-1273-induced IgG mediates protection against SARS-CoV-2

High titer antibody responses in blood and upper and lower airways associated with the rapid control of viral load and lower airway pathology in the lung suggested that antibody was the primary immunological mechanism of protection. To directly address whether vaccine-induced antibody was sufficient to mediate protection, mRNA-immune NHP IgG was purified from pooled sera 2 weeks post-boost of 100 μg of mRNA-1273 (*13*) (Fig. 5A) and passively transferred to hamsters (Fig. 5B). Two or 10 mg of total mRNA-1273-immune NHP IgG or 10 mg of pre-immune NHP IgG (control) was administered to 8 Syrian hamsters/group, and immediately before challenge, humoral S-specific IgG (Fig. 5C) and pseudovirus neutralization titers (Fig. 5D) were assessed to confirm the relative antibody responses prior to the passive transfer. Of note, while there were different antibody responses based on the two different amounts of total IgG transferred, this study is only used for showing antibody is sufficient for protection and not for making any determinations of correlates. At 24 hr post-immunization, hamsters were inoculated with 3 × 10^4^ PFU SARS-CoV-2 (USA-WA1/2020) intranasally. Control hamsters and hamsters that received 2 mg mRNA-1273-immune IgG showed a ~10% weight loss by day 6, a defined endpoint for pathogenicity in this model (*32*). By contrast, hamsters that received 10 mg mRNA-1273-immune NHP IgG showed little to no weight loss post-challenge (Fig. 5E). These data show that mRNA-1273 immune IgG is sufficient to mediate protection from disease *in vivo* against SARS-CoV-2 infection.

**Fig. 5.**
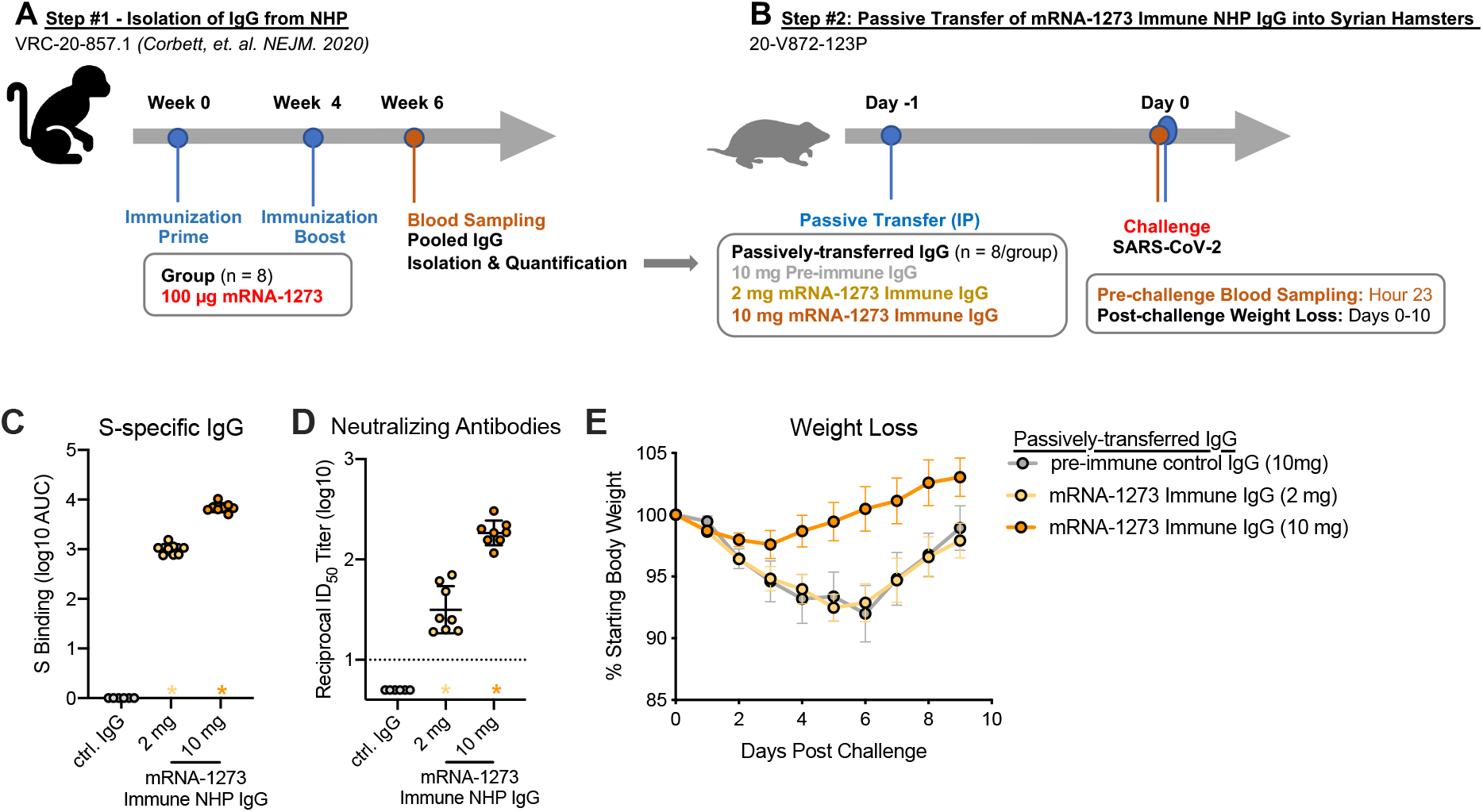
Passive transfer of mRNA-1273 immune NHP IgG into Syrian hamsters. (A) Sera were pooled from all NHP that received 100 μg of mRNA-1273 in a primary vaccination series. (B) mRNA-1273 immune NHP IgG (2 mg, yellow or 10 mg, orange) or pre-immune NHP IgG (10 mg, gray) was passively transferred to Syrian hamsters (n = 8/group) 24 hours prior to SARS-CoV-2 challenge. Twenty-three hours post-immunization, hamsters were bled to quantify circulating S-specific IgG (C) and SARS-CoV-2 pseudovirus neutralizing antibodies (D). Following challenge, hamsters were monitored for weight loss (E). (C-D) Circles represent individual NHP. Bars and error bars represent GMT and geometric SD, respectively. Asterisks at the axis represent animals that did not receive adequate IgG via passive transfer and were thus excluded from weight loss analyses. (D) The dotted line indicates the neutralization assay limit of detection. (E) Circle and error bars represent mean and SEM, respectively.

## Discussion

Defining immune correlates of protection is a critical aspect of vaccine development for extending the use of approved vaccines and facilitating the development of new candidate vaccines, as well as defining potential mechanisms of protection. For SARS-CoV-2, a primary goal of current vaccines is to prevent symptomatic COVID-19. For moderate to severe disease this is likely a consequence of reducing viral load in lower airways and for mild disease reducing viral load in both lower and upper airways. An additional benefit of upper airway protection is that limiting nasal carriage of virus will also reduce transmission risk. Here, we establish that the level of S-specific antibody elicited by mRNA-1273 vaccination correlates with control of upper and lower airway viral replication following SARS-CoV-2 challenge in NHP. Furthermore, we find that CD4 T cell responses elicited by the vaccine did not provide any additional power to predict protection but were associated with reduction of viral load in NS. Finally, we show that vaccine-elicited antibodies are sufficient for protection against disease in hamsters. S-binding antibodies met our prespecified statistical analysis criteria for a correlate of protection as they strongly inversely correlated with lower and upper airway viral loads with no additional predictive power provided by vaccine dose.

A key parameter to assess correlates of protection in NHP is the amount of virus used for challenge. In this study, a challenge dose of 8×10^5^ PFU of a newly characterized, pathogenic SARS-CoV-2 USA-WA1/2020 strain was used to achieve a level of viral replication comparable to or exceeding that observed from nasal swabs of humans with symptomatic infection, measured by sgRNA (*33*) or genomic vRNA (*34–36*). The sgRNA levels for N or E of 10^6^-10^7^ in control animals at day 2 post-challenge are among the highest reported for NHP challenge studies and likely models viral load in humans at the upper end of inoculum size. We used the same qualified antibody binding and pseudovirus neutralization assays for assessing immune responses as in human Phase 3 vaccine trials. Additionally, the use of WHO standards to report binding titers in IU enables comparison of immune responses and outcomes with other NHP vaccine studies and benchmarking to human vaccine clinical trials. We show that a 10-fold increase in S-binding titers was associated with approximately 10-fold reductions in viral replication in BAL and NS post-challenge. No animal with S-specific IgG >336 IU/mL had BAL sgRNA >10,000 copies/mL, and no animal with S-specific IgG >645 IU/mL had NS sgRNA >100,000 copies/swab, which were chosen as thresholds for protection. These reductions in viral replication, assessed by sgRNA compared to controls, were associated with limited inflammation and viral antigen detection in the lung tissue and may be sufficient to prevent moderate or severe lower airway infection. Notably, even animals in the 1 and 3 μg dose groups, for which the elicited S-specific antibody levels were 81 and 272 IU/mL and reciprocal pseudovirus neutralization titers of 49 and 53, exhibited ~2-4 log_10_ less viral replication in BAL compared to the control animals at day 2 post-challenge. Last, S-binding titers of >488 IU/mL were associated with no increase in N-specific antibody responses post-challenge, a metric used in human studies for assessing prevention of asymptomatic infection post-vaccination (*34*).

The antibody titers required in this high-dose challenge NHP model for a reduction in viral replication may be a conservative estimate for what is required to prevent clinical disease in humans. The strong correlation and proportional changes between S-specific binding titers with serum neutralizing activity support using easy-to-measure binding titers as the primary metric for defining a correlate of protection in humans at least for mRNA-based vaccines delivering similar antigens and eliciting similar patterns of immunogenicity.

Mucosal antibody responses are thought to be an important mechanism of protection against a variety of upper respiratory viral infections (*37–41*). Both BAL and nasal wash S-specific IgG and IgA were predictive for reducing sgRNA in these compartments. Serum antibody levels were a strong predictor of IgG and IgA responses in BAL and nasal washes as well as for protection measured by viral replication in these sites. Given mRNA-1273 is administered via intramuscular delivery, these data suggest that localized upper and lower airway antibodies are transudated from serum and suggest serum antibody levels to be a surrogate for BAL and NS antibody levels following mRNA-1273 vaccination.

Additionally, here we considered whether the infection boosted vaccine-induced antibodies. There were no anamnestic S-specific responses or increase in N-specific responses in blood or BAL within 28 days post infection in the >3 μg dose groups compared to pre-challenge, consistent with these doses inducing higher antibody responses. By contrast, there were increased anamnestic S-binding antibody responses in the <1 μg dose groups. These data suggest that boosting of vaccine-induced antibodies can occur following upper airway infection in animals that have minimal viral replication in the lower airway. Therefore, the necessity and timing of subsequent vaccine boosting will depend on whether the goal is to prevent severe disease and lower airway infection while allowing community exposure to provide mucosal immunity from upper airway infection or to achieve sterilizing immunity through vaccination to more rapidly reduce transmission.

Based on the rapid control of viral replication in the lower airway by day 2 post-challenge and the presence of robust airway IgG responses, we hypothesized that antibodies are not only a correlate but also the primary mechanism of protection. The critical role of antibodies for mediating protection is also consistent with multiple human cohort studies assessing antibody responses in people after prior exposure showing that subsequent infection is reduced (*42–45*). The level of serum neutralizing activity has also been suggested as a predictor of efficacy from COVID-19 (*45*). The demonstration here that purified mRNA-1273 immune NHP IgG is associated with dose-dependent protection from weight loss in the highly pathogenic hamster model provides direct evidence that vaccine-elicited antibody is sufficient to mediate protection from disease.

To conclude, this study establishes a critical role of functional antibodies as a correlate of protection against SARS-CoV-2 in a NHP and that for mRNA expressing this particular spike antigen, binding antibody is a surrogate marker of protection. Furthermore, this study establishes that a lower antibody level is needed for reduction of viral replication in the lower airway than in the upper airway. Ongoing NHP studies will assess durability of mRNA-1273-elicited protection and efficacy of mRNA-1273 vaccination against global SARS-CoV-2 variants. These findings anticipate the correlates analysis comparing virus replication in NS with serum antibody being performed on samples from vaccinated subjects in Phase 3 clinical trials who experienced breakthrough infection.

## Acknowledgements

We thank Tracy Ruckwardt, Nicole Doria-Rose, and additional members of all included laboratories for critical discussions and advice pertaining to experiments included in the manuscript. We thank Judy Stein and Monique Young for technology transfer and administrative support, respectively. We thank members of the NIH NIAID VRC Translational Research Program, including Chris Case, Hana Bao, Elizabeth McCarthy, Jay Noor, Alida Taylor, and Ruth Woodward, for technical and administrative assistance with animal experiments. We thank Huihui Mu and Michael Farzan for the ACE2-overexpressing 293 cells. We thank the laboratory of Peter Kwong for providing protein for use in ELISA assays for detection of mucosal antibodies. We thank Andy Pekosz for the B.1.351 variant used in FRNT assays and Eli Boritz for assistance with B.1.351 sequencing and analysis. We thank Michael Brunner and Dr. Michael Whitt for kind support on recombinant VSV-based SARS-CoV-2 pseudovirus production.

## Funding

Intramural Research Program of the VRC, NIAID, NIH

Department of Health and Human Services, Office of the Assistant Secretary for Preparedness and Response, Biomedical Advanced Research and Development Authority, Contract 75A50120C00034

Undergraduate Scholarship Program, Office of Intramural Training and Education, Office of the Director, NIH (K.S.C.)

NIAID Research Participation Program, administered by the Oak Ridge Institute for Science and Education through an interagency agreement between the U.S. Department of Energy and NIAID (R.W.)

Emory Executive Vice President for Health Affairs Synergy Fund Award (M.S.S.)

Pediatric Research Alliance Center for Childhood Infections and Vaccines and Children’s Healthcare of Atlanta (M.S.S.)

Woodruff Health Sciences Center 2020 COVID-19 CURE Award (M.S.S.)

## Author Contributions

K.S.C., M.C.N, B.F., M.G., S.O., T.S.J., S.N.S., V.V.E., K.F., L.L., C.M., J.F., B.F., K.W., A.C., M.K., A.P.W., J.I.M., O.M.A., S.F.A., M.M.D., J.F., D.R.F., E.L., A.T.N., S.T.N., S.J.P., A.C., A.D., A.F., J.G., S.K., L.P., M.P., K.S., D.V., S.Z., K.W.B., M.M., B.M.N., R.V., H.A., K.E.F., D.K.E., J.R.M., I.N.M., M.G.L., A.C., D.M., M.S.S, A.M., N.J.S., M.R., D.C.D., B.S.G., and R.A.S. designed, completed, and/or analyzed experiments. O.M.A., S.B-B., K.L., W.S., E.S.Y., Y.Z., and L.W. provided critical published reagents/analytic tools. K.S.C., M.C.N., N.J.S., M.R., B.S.G, and R.A.S. wrote the manuscript. K.S.C., M.C.N. M.G., and G.A. prepared figures and tables. All authors contributed to discussions about and editing of the manuscript.

## Data and materials availability

All data are available in the main text or the supplementary materials.

