## Supplementary Material for "Immune Correlates of Protection by mRNA-1273 Immunization against SARS-CoV-2 Infection in Nonhuman Primates"

**This PDF file includes:**

Materials and Methods

Figs. S1 to S10

Tables S1 to S2

**Materials and Methods**

Preclinical mRNA-1273 mRNA and Lipid Nanoparticle Production

A sequence-optimized mRNA encoding prefusion-stabilized SARS-CoV-2 S-2P protein (*2, 3*) was synthesized *in vitro* and formulated as previously reported (*13, 25*).

Rhesus Macaque Model

Animal experiments were carried out in compliance with all pertinent US National Institutes of Health regulations and approval from the Animal Care and Use Committee of the Vaccine Research Center and Bioqual, Inc. (Rockville, MD). Studies were conducted at Bioqual, Inc. The experimental details of VRC-20-857.1 (Fig. S1A) are previously published (*13*). For VRC-20-857.3 (Fig. S1B) or VRC-20-857.4 (Fig. S1C), 3–8-year-old male Indian-origin rhesus macaques were sorted by age and weight and then stratified into groups. Animals were immunized intramuscularly (IM) at week 0 and week 4 with doses ranging from 0.3 - 30 μg of mRNA-1273 in 1 mL of 1X PBS into the right hindleg. Placebo-control animals were administered equal volume of 1X PBS. At week 8 (4 weeks post-boost), all animals were challenged with a total dose of $8x10^5$ PFU SARS-CoV-2. The stock of $1.99x10^6$ TCID50 or $3x10^6$ PFU/mL SARS-CoV-2 USA-WA1/2020 strain (BEI: NR-70038893) was diluted and administered in 3 mL by the intratracheal route and 1 mL by the intranasal route (0.5 mL per nostril). Pre- and post-challenge sample collection is detailed in Fig. S1C.

Passive Transfer of Purified IgG into Golden Syrian Hamster Model

Sera from NHP immunized with 100 µg of mRNA-1273 in VRC-20-857.1 (*13*) (Fig. S1A) were collected 2 weeks post-boost and pooled. Total IgG was purified from pooled sera using Protein G Sepharose 4 Fast Flow resin (Cytiva), according to manufacturer's instructions, and quantified by NanoDrop™ OneC Microvolume UV-Vis Spectrophotometer (Thermo Scientific). The eluted protein dialyzed against 1X PBS pH7.4 (GIBCO) and concentrated to 10 µg/ml using Amicon® Ultra centrifugal Filter (Millipore Sigma). Golden Syrian hamsters, aged 6-8 weeks old, were randomized into groups of 8 based on weight, with each group containing a 1:1 male:female ratio. IgG was passively transferred by intraperitoneal (IP) injection 1 day prior to challenge. Immediately prior to challenge, sera were collected for antibody assessments. Hamsters were inoculated intranasally with $3\times10^4$ PFU USA-WA1/2020 SARS-CoV-2 (BEI, NR-53780) in a final volume of 100 µL and split between each nostril. Body weight and clinical observations were made daily post-challenge.

Quantification of SARS-CoV-2 sgRNA

BAL and NS were collected 2-, 4-, and 7-days post-challenge. At the time of collection, NS were frozen in 1 mL of 1X PBS containing 1 µL of SUPERase-In RNase Inhibitor (Invitrogen) and frozen at -80°C until extraction. Nasal specimens were thawed at 55°C, and the swab removed. The remaining PBS was mixed with 2 mL of RNAzol BD (Molecular Research Center) and 20 µL acetic acid. At the time of collection, 1 mL of BAL fluid was mixed with 1 mL of RNAzol BD containing 10 µL acetic acid and frozen at -80°C until extraction. BAL specimens were thawed at room temperature (RT) and mixed with an additional 1 mL of RNAzol BD containing 10 μL acetic acid.

Total RNA was extracted from nasal specimens and BAL fluid using RNAzol BD Column Kits and eluted in 65 μL water. Subgenomic SARS-CoV-2 E and N mRNA was quantified via reverse transcription-polymerase chain reaction (PCR) using a technique similar to that described previously (*33*). Reactions were conducted with 5 μL RNA and TaqMan Fast Virus 1-Step Master Mix (Applied Biosystems) with 500 nM primers and 200 nM probes. Primers and probes were as follows: sgLeadSARSCoV2_F: 5'-CGATCTCTTGTAGATCTGTTCTC-3', E subgenomic mRNA - E_Sarbeco_P: 5'-FAM-ACACTAGCCATCCTTACTGCGCTTCG-BHQ1-3' and E_Sarbeco_R: 5'-ATATTGCAGCAGTACGCACACA-3', N subgenomic mRNA - wtN_P: 5'-FAM-TAACCAGAATGGAGAACGCAGTGGG-BHQ1-3'and wtN_R: 5'-GGTGAACCAAGACGCAGTAT-3'.

Reactions were run on a QuantStudio 6 Pro Real-Time PCR System (Applied Biosystems) at the following conditions: 50°C for 5 min, 95°C for 20 sec, and 40 cycles of 95°C for 15 sec and 60°C for 1 min. Absolute quantification was performed in comparison to a standard curve. For the standard curve, the E or N subgenomic mRNA sequence was inserted into a pcDNA3.1 vector (Genscript) and transcribed using MEGAscript T7 Transcription Kit (Invitrogen) followed by MEGAclear Transcription Clean-Up Kit (Invitrogen). The lower limit of quantification was 50 copies.

Histopathology and Immunohistochemistry (IHC)

As previously described (*13*), NHP lung tissue sections were stained with hematoxylin and eosin (H&E) for routine histopathology and a rabbit polyclonal SARS-CoV-2 (GeneTex, GTX135357) for detection of SARS-CoV-2 virus antigen. All samples were evaluated by a boarded-certified veterinary pathologist.

4-Plex Meso-Scale ELISA

MSD SECTOR® plates are precoated by MSD with SARS-CoV proteins (S-2P, RBD, and N) and a Bovine Serum Albumin (BSA) control in each well in a specific spot-designation for each antigen. The assay will be performed with a Beckman Coulter Biomek based automation integration platform including the Biotek 405TS Plate Washer. Serum samples will be heat-inactivated for 30 min at 56°C prior to assay. Plates are blocked for 60 min at RT with MSD blocker A solution without shaking. Plates are washed and MSD reference standard (calibrator), QC test sample (pool of COVID-19 convalescent sera) and human serum test samples are added to the precoated wells in duplicates in an 8-point dilution series. Reference standard is added in triplicates. MSD control sera (low, medium and high) are added undiluted in triplicates as per validated assay format. Additional assay controls might be added in triplicates. Samples are incubated at RT for 4 hr with shaking on a Titramax Plate shaker (Heidolph) at 1500 rpm. SARS-CoV-2 specific antibodies present in the sera or controls bind to the coated antigens. Plates are washed to remove unbound antibodies. Antibodies bound to the SARS-CoV-2 viral proteins are detected using an MSD SULFO-TAG™ anti-human IgG detection antibody incubated for 60 min at RT and with shaking. Plates are washed and a read solution (MSD GOLD™ read buffer) containing electrochemiluminescence (ECL) substrate is applied to the wells, and the plate is entered into the MSD MESO Sector S 600 detection system. An electric current is applied to the plates and areas of well surface which form antigen-anti human IgG antibody SULFO-TAG™ complex will emit light in the presence of the ECL substrate.

The MSD MESO Sector S 600 detection system quantitates the amount of light emitted and reports the ECL unit response as a result for each test sample, control sample and reference standard of each plate. Analysis is performed with the MSD Discovery Workbench software, Version 4.0. Calculated ECLIA parameters to measure binding antibody activities will include interpolated concentrations or assigned arbitrary units (AU/mL) read from the standard curve. Recently the arbitrary units were bridged to the WHO International Standard and a conversion factor was calculated and confirmed. Parallelism was established for all three antigens between the MSD provided reference standard and the WHO provided international standard. Concentration assignments were performed and then confirmed both at MSD and as part of a multi-site confirmation study. Sample results reported here have been converted to international units (IU/mL).

ELISA for Temporal NHP Serum Antibodies and Hamster Serum Antibodies

SARS-CoV-2 S-specific IgG in serum was quantified by ELISA; the methods used were similar to those previously published (*25*). Here, the only amendment to previously published methodology is the resulting data are depicted as endpoint titers, which were calculated as the mean serum titer that reached 10X SD using Prism v9.0.2 (GraphPad).

Meso-Scale ELISA for Mucosal Antibody Responses

Using previously described methods (*46*), total S-specific IgG and IgA were determined by MULTI-ARRAY ELISA using Meso Scale technology (Meso Scale Discovery, MSD).

Serum Antibody Avidity Assay

Avidity was assessed using an ammonium thiocyanate (NaSCN)-based avidity ELISA against the full-length SARS-CoV-2 S-2P antigen. ELISA was performed as described previously in the absence or presence of chaotropic agent. Briefly, plasma/serum samples were serially diluted in blocking buffer and 100 mL was transferred to the plates. After 1-hr incubation, plates were washed and half of the samples were then incubated with 100 mL PBS while the other half of the paired samples were treated with 100 mL of 0.1-1.0 M sodium thiocyanate solution (NaSCN, Sigma-Aldrich) for 15-30 min at RT followed by washing, and incubated for 1 hr with 100 mL of goat-anti-human IgG (H+L, Cat#PA-1-8463) or goat-anti-monkey IgG (H+L, Cat#A18811) secondary antibody conjugated to horseradish peroxidase (HRP, Thermo Fisher) with detection using SureBlueTM TMB Microwell Peroxidase Substrate (1-Component, SeraCare, Cat#52-00-01). The avidity index was calculated using the ratio of IgG binding to S-2P in the absence or presence of NaSCN. The reported AI is the average of two independent experiments, each containing duplicate samples.

ACE2 Binding Inhibition Assay

ACE2 binding inhibition was completed, as previously described (*47*), on 1:40 diluted sera samples using Mesoscale Discovery 384-well, 4-Spot Custom Serology SECTOR® plates pre-coated with SARS-CoV-2 RBD. Binding was detected using SULFO-TAG™ labeled ACE2. Both reagents were generously supplied by the manufacturer.

Lentiviral Pseudovirus Neutralization Assay

As previously described (*48*), pseudotyped lentiviral reporter viruses were produced by the co-transfection of plasmids encoding S proteins from Wuhan-1 strain (Genbank #: MN908947.3) with a D614G mutation, a luciferase reporter, lentivirus backbone, and human transmembrane protease serine 2 (TMPRSS2) genes into HEK293T/17 cells (ATCC CRL-11268). Sera, in duplicate, were tested for neutralizing activity against the D614G pseudoviruses by quantification of luciferase activity [in relative light units (RLU)]. Percent neutralization was normalized considering uninfected cells as 100% neutralization and cells infected with pseudovirus alone as 0% neutralization. $IC_{50}$ titers were determined using a log(agonist) vs. normalized-response (variable slope) nonlinear regression model in Prism v9.0.2 (GraphPad).

VSV Pseudovirus Neutralization Assay

To make SARS-CoV-2 pseudotyped recombinant VSV-ΔG-firefly luciferase virus, BHK21/WI-2 cells (Kerafast, EH1011) were transfected with the S plasmid expressing full-length S with D614G mutation and subsequently infected with VSVΔG-firefly-luciferase as previously described (*49*). Neutralization assays were completed on A549-ACE2-TMPRSS2 cells with serially diluted serum samples as previously described (*16*).

Focus Reduction Neutralization Test (FRNT)

VeroE6 cells were obtained from ATCC (clone E6, ATCC, #CRL-1586) and cultured in complete DMEM medium consisting of 1X DMEM (VWR, #45000-304), 10% FBS, 25mM HEPES Buffer (Corning Cellgro), 2 mM L-glutamine, 1 mM sodium pyruvate, 1X Non-essential Amino Acids, and 1X antibiotics. EHC-083E (D614G SARS-CoV-2) was previously described (*50*). Viruses were propagated in Vero-TMPRSS2 cells to generate viral stocks. Viral titers were determined by focus-forming assay on VeroE6 cells. Viral stocks were stored at -80°C until use. FRNT assays were performed as previously described (*51*). Briefly, samples were diluted at 3-fold in 8 serial dilutions using DMEM (VWR, #45000-304) in duplicates with an initial dilution of 1:10 in a total volume of 60 ml. Serially diluted samples were incubated with an equal volume of SARS-CoV-2 (100-200 foci per well) at 37° C for 1 hr in a round-bottomed 96-well culture plate. The antibody-virus mixture was then added to Vero cells and incubated at 37° C for 1 hr. Post-incubation, the antibody-virus mixture was removed and 100 μl of prewarmed 0.85% methylcellulose (Sigma-Aldrich, #M0512-250G) overlay was added to each well. Plates were incubated at 37° C for 24 hr. After 24 hr, methylcellulose overlay was removed, and cells were washed three times with PBS. Cells were then fixed with 2% paraformaldehyde in PBS (Electron Microscopy Sciences) for 30 min. Following fixation, plates were washed twice with PBS and 100 µl of permeabilization buffer (0.1% BSA [VWR, #0332], Saponin [Sigma, 47036-250G-F] in PBS), was added to the fixed Vero cells for 20 min. Cells were incubated with an anti-SARS-CoV S primary antibody directly conjugated to biotin (CR3022-biotin) for 1 hr at RT. Next, the cells were washed 3x in PBS and avidin-HRP was added for 1 hr at RT followed by three washes in PBS. Foci were visualized using TrueBlue HRP substrate (KPL, # 5510-0050) and imaged on an ELISPOT reader (CTL). Antibody neutralization was quantified by counting the number of foci for each sample using the Viridot program (*52*). The neutralization titers were calculated as follows: 1 - (ratio of the mean number of foci in the presence of sera and foci at the highest dilution of respective sera sample). Each specimen was tested in duplicate. The FRNT-50 titers were interpolated using a 4-parameter nonlinear regression in GraphPad Prism v9.0.2.4.3. Samples that do not neutralize at the limit of detection at 50% are plotted at 5 and was used for geometric mean calculations.

Intracellular Cytokine Staining

Cryopreserved peripheral-blood mononuclear cells were thawed, rested overnight, and stimulated with SARS-CoV-2 S protein (S1 and S2, homologous to the vaccine insert) and costimulatory antibodies anti-CD28 and anti-CD49d (clones CD28.2 and 9F10, BD Biosciences). Negative controls received an equal concentration of dimethyl sulfoxide (without peptides) and costimulatory antibodies. Cytokine staining was performed as described previously (*13*).

Correlations and Statistical Analysis

Graphs show data from individual animals with dotted lines indicating assay limits of detection. Correlations between antibody measurements, and between antibody measurements and dose of vaccine, are estimated and tested against 0 using Spearman's nonparametric correlation. Univariate and multivariate linear models are used to evaluate the potential role of antibodies as correlates of protection; following the Prentice criteria (*53*), we categorize antibody measures as potential correlates of protection if they are significantly univariately predictive of $\log_{10}$ sgRNA as a measure of viral replication, and if the addition of $\log_{10}$ dose to the linear model does not significantly improve prediction, as assessed by a likelihood ratio test. Univariate and multivariate linear models are also used to explore the combination of t-cell responses and antibody responses in prediction of sgRNA. Comparisons between pre-challenge and post-challenge were assessed using paired t-tests, based on the last measurement (day 28 post-challenge when available, day 14 when not). Intracellular cytokines likely response labels were derived using the MIMOSA[(*54*)] package. Analyses were done in R version 4.0.2 and Prism v9.0.2.

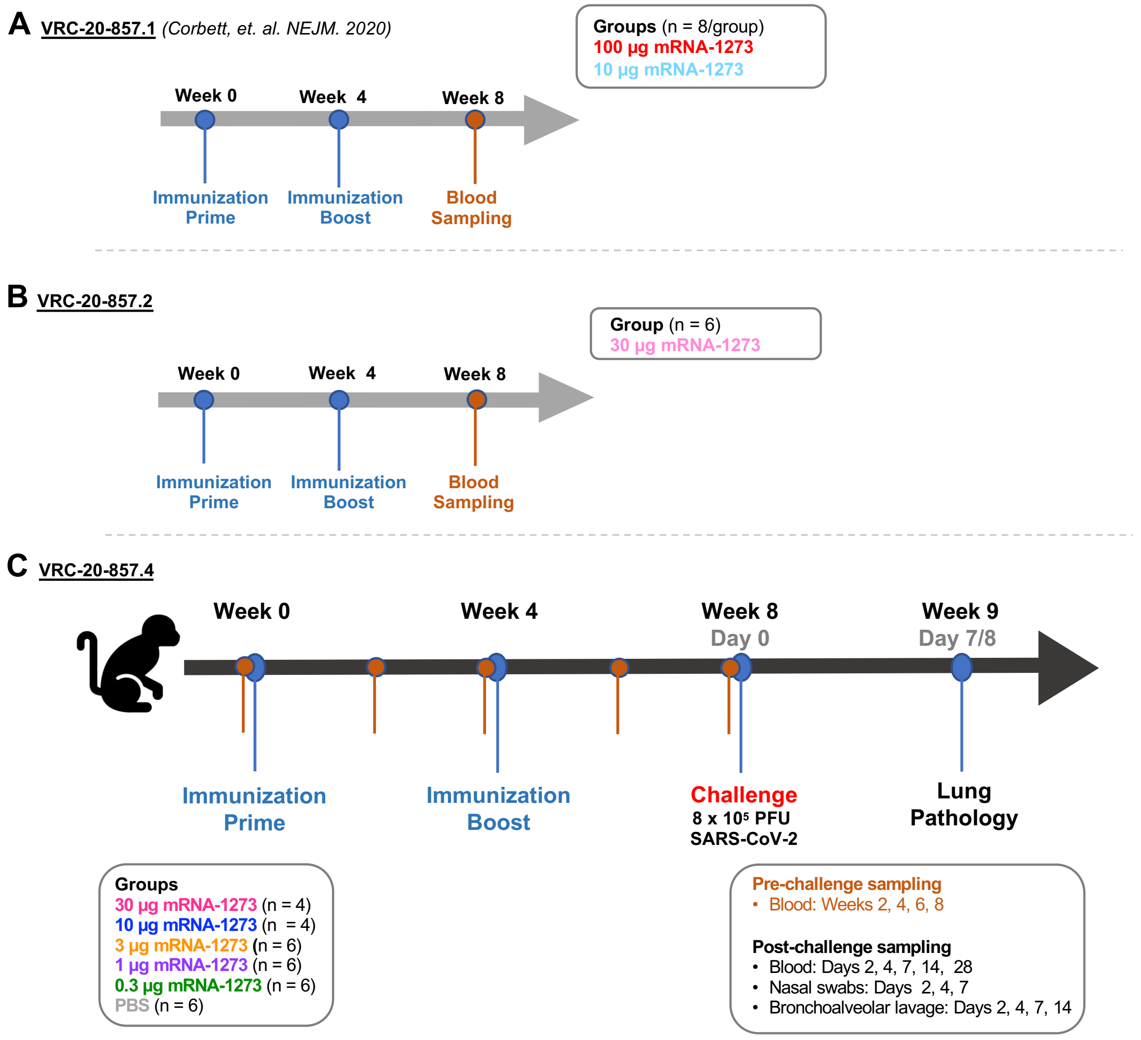


**Fig. S1. Study design: Evaluation of immune correlates of protection of mRNA-1273 in rhesus macaques.** (A-B) The design of the correlates of protection study (C) was informed by previous studies to assess the immunogenicity of various doses of mRNA-1273 in NHP. (C) To assess immune correlates of protection, Rhesus macaques were immunized at 0 and 4 weeks with PBS or various doses of mRNA-1273 and challenged 4 weeks post-boost with a total of 8 x $10^5$ PFU of SARS-CoV-2. The viral inoculum was administered as 6 x $10^5$ PFU in 3 mL intratracheally and 2 x $10^5$ PFU in 1 mL intranasally (0.5 mL into each nostril). Sera were collected preimmunization and bi-weekly post-prime and post-boost. Sera, nasal washes, and bronchoalveolar lavages were collected post-challenge on days 2, 4, 7, 14, and 28. Lung pathology was assessed on days 7 and 8 post-challenge in a subset of animals (n = 1 or 2/group). Pre-challenge serological assessments are overlayed within applicable Fig.s where squares represent experiments VRC-20-857.1 (A) and VRC-20-857.2 (B), and circles represent experiment VRC-20-857.4 (C).

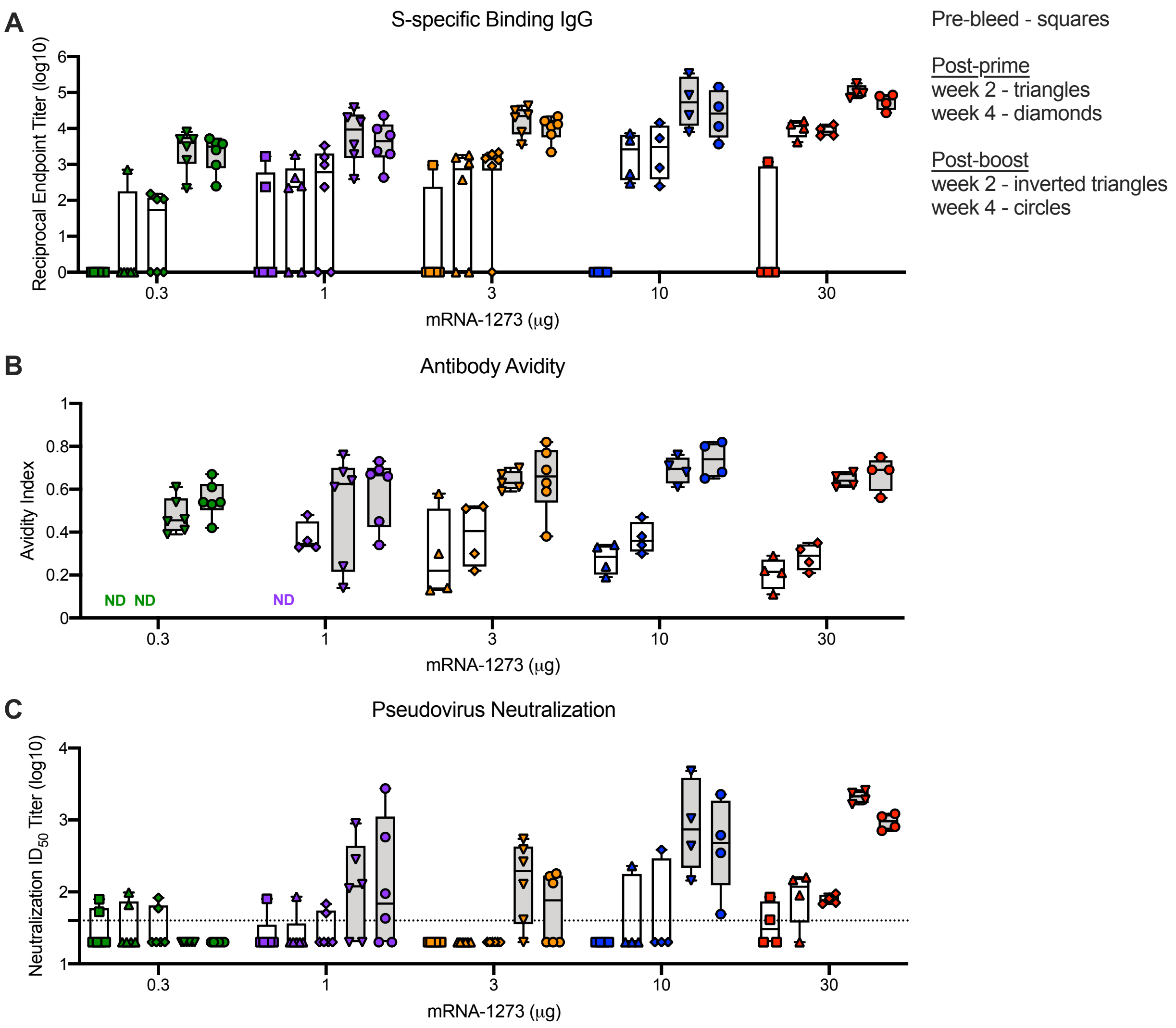


**<u>Fig. S2</u>. Temporal serum antibody responses following mRNA-1273 immunization.** Rhesus macaques were immunized according to Fig. S1C. Sera collected at week 0 (pre-bleed, unfilled bars) and 2- and 4-weeks post-prime (unfilled bars) and post-boost (filled bars) and subsequently assessed for SARS-CoV-2 S-specific IgG (A), antibody avidity (B), and SARS-CoV-2 D614G lentiviral-based pseudovirus neutralization (C). Symbols represent individual NHPs. Boxes and horizontal bars denote interquartile ranges (IQR) and medians, respectively; whisker end points are equal to the maximum and minimum values. Dotted lines indicate assay limit of detection, where applicable. ND = not determined due to low-level S-specific binding antibody titers.

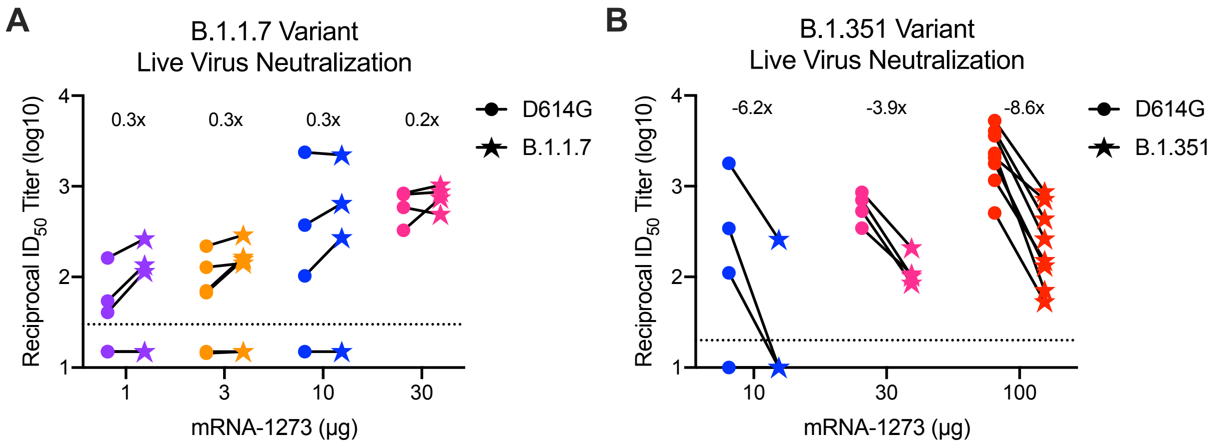


**Fig. S3. Ability of mRNA-1273 immune NHP serum to neutralize global variants.** Rhesus macaques were immunized according to Fig. S1. Sera collected 4-weeks post-boost, were assessed for focus reduction neutralization using D614G-encompassing SARS-CoV-2 EHC-83E compared to B.1.1.7 (A) and B.1.351 (B) SARS-CoV-2 variants. Symbols represent individual NHP. For each immunogen group, geometric mean titer (GMT) for each variant was compared to D614G, and fold-change is indicated above respective before and after plots.

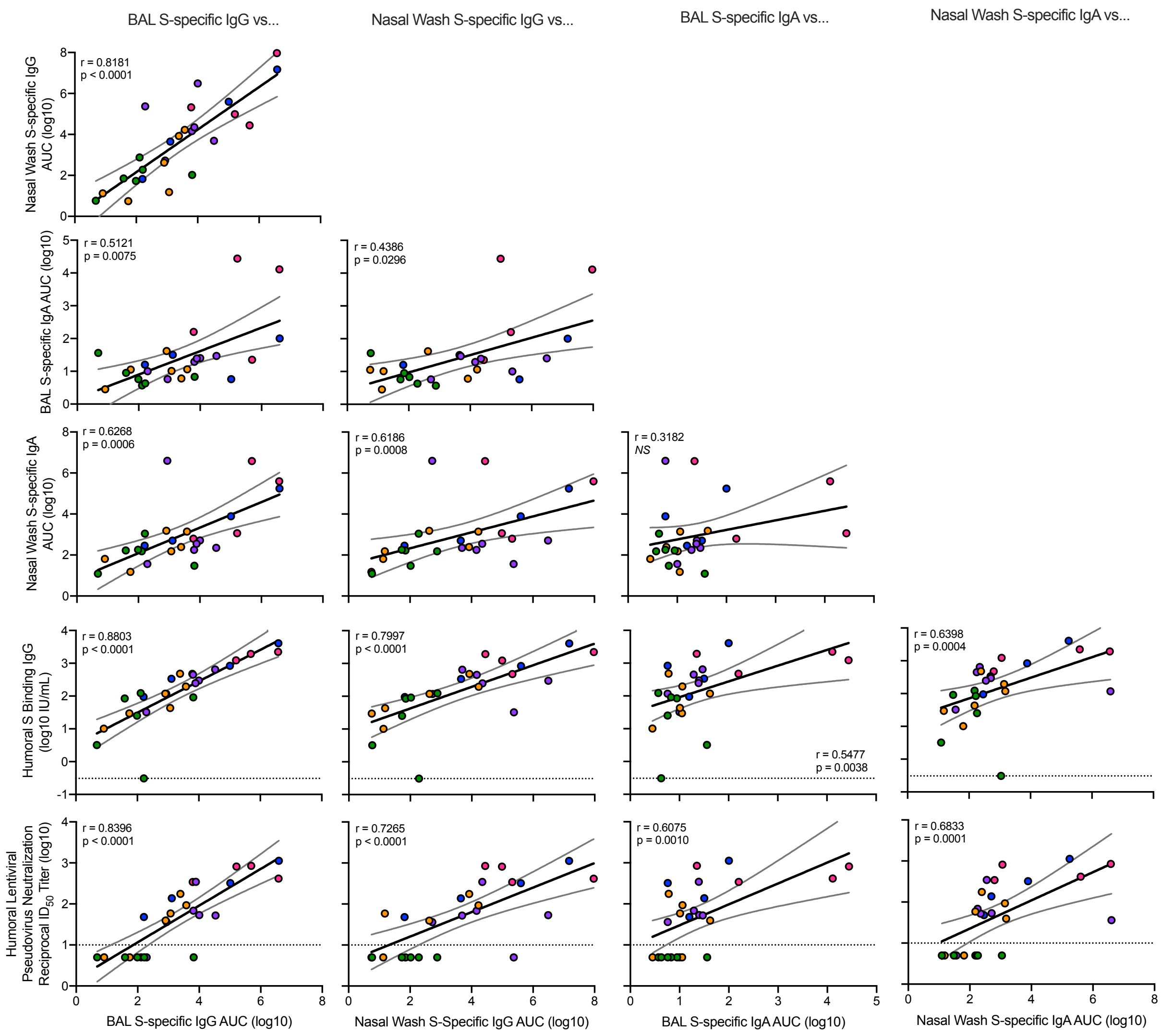


**<u>Fig. S4</u>. Correlations of mucosal antibody responses to humoral antibody responses.** Rhesus macaques were immunized according to Fig. S1C. Plots show correlations between S-specific BAL IgG, nasal wash IgG, BAL IgA, and nasal wash IgA at week 2 post-boost with each other and with humoral S-specific IgG and lentiviral-based pseudovirus neutralization at week 4 post-boost. Circles represent individual NHP, where colors indicate mRNA-1273 dose. Dotted lines indicate assay limits of detection. Black and gray lines indicate linear regression and 95% confidence interval, respectively. 'r' represents Spearman's correlation coefficients, and 'p' the corresponding p-values.

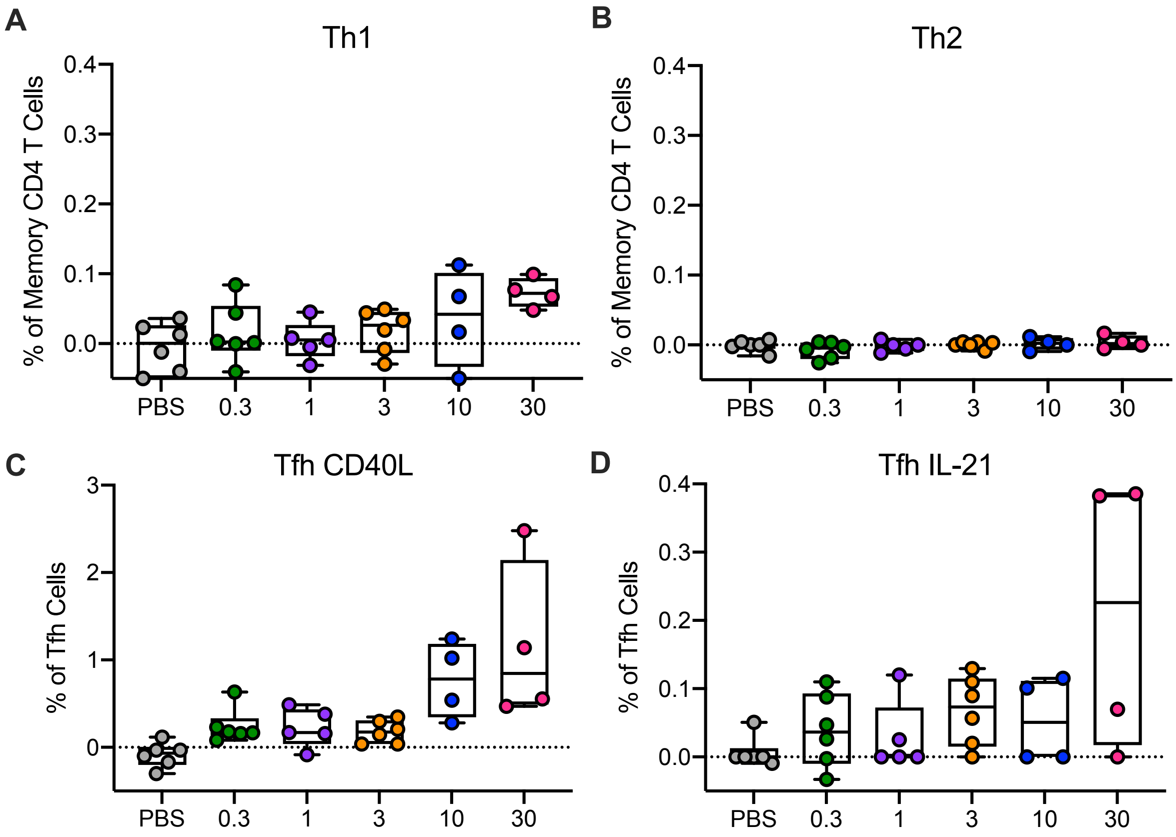


**<u>Fig. S5</u>. T cell responses following mRNA-1273 immunization.** Rhesus macaques were immunized according to Fig. S1C. Intracellular staining was performed on PBMCs at week 6, 2 weeks following boost, to assess T cell responses to SARS-CoV-2 S protein peptide pools, S1 and S2. Responses to S1 and S2 individual peptide pools were summed. (A) Th1 responses (IFNg, IL-2, or TNF), (B) Th2 responses (IL-4 or IL-13), (C) Tfh CD40L upregulation (peripheral follicular helper T cells (Tfh) were gated on central memory $CXCR5^{+}PD\text{-}1^{+}ICOS^{+}$ CD4 T cells), (D) Tfh IL-21. Boxes and horizontal bars denote IQR and medians, respectively; whisker end points are equal to the maximum and minimum values. Circles represent individual NHP. Dotted lines are set to 0%.

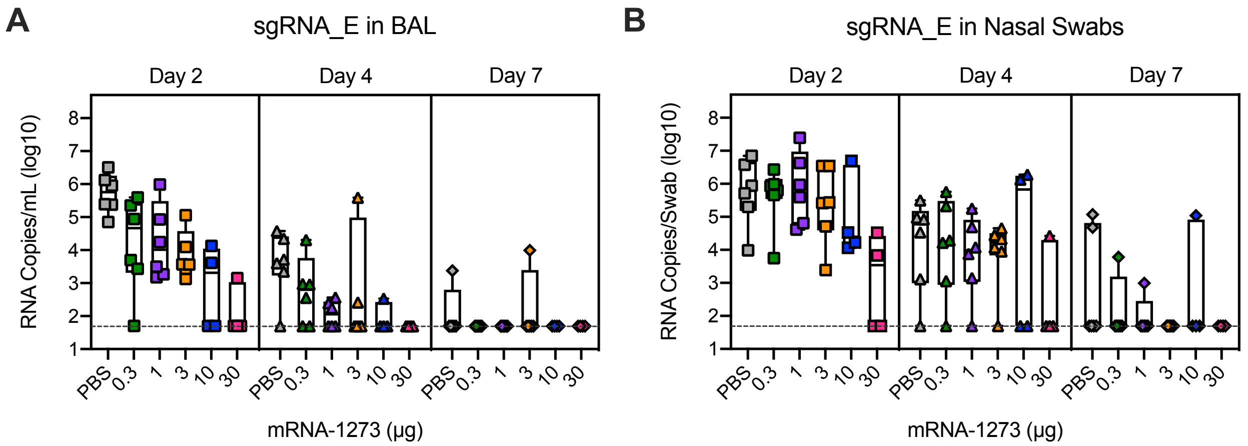


**<u>Fig. S6</u>. Efficacy of mRNA-1273 against upper and lower respiratory viral replication.** Rhesus macaques were immunized and challenged as described in Fig. S1C. BAL (A) and NS (B) were collected on days 2 (squares), 4 (triangles), and 7 (diamonds) post-challenge, and viral replication was assessed by detection of SARS-CoV-2 E-specific sgRNA. Boxes and horizontal bars denote the IQR and medians, respectively; whisker end points are equal to the maximum and minimum values. Symbols represent individual NHP and overlap for equal values where constrained. Dotted lines indicate assay limits of detection.

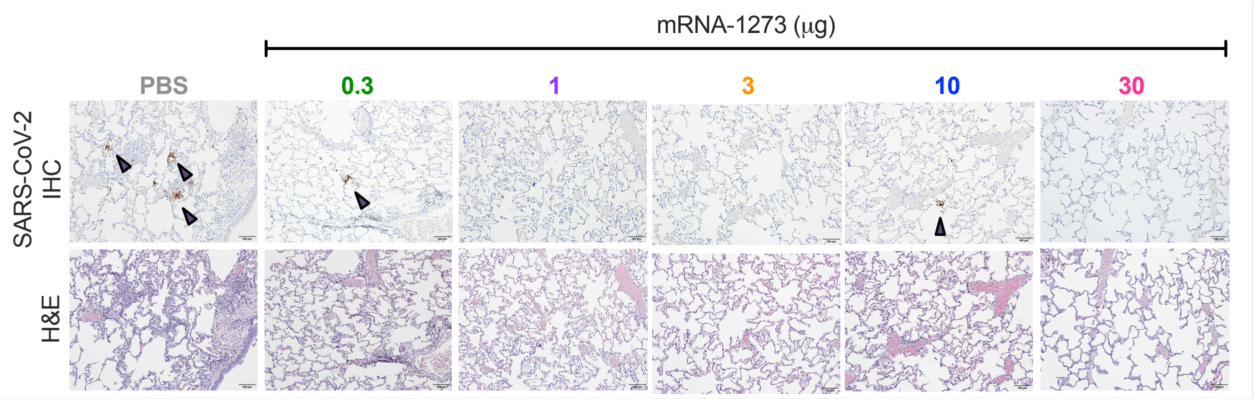


**<u>Fig. S7</u>. Post-challenge lung histopathological analysis and viral detection.** Rhesus macaques were immunized and challenged as described in Fig. S1C. Seven days after challenge, lung samples (n = 1/group) were evaluated for evidence of virus infection (top) and the presence of inflammation (bottom); representative images show the location and distribution of SARS-CoV-2 virus antigen (IHC) in serial lung tissue sections. Arrows indicate areas positive for viral antigen. Each image is taken at 10x magnification; scale bars represent 100 microns.

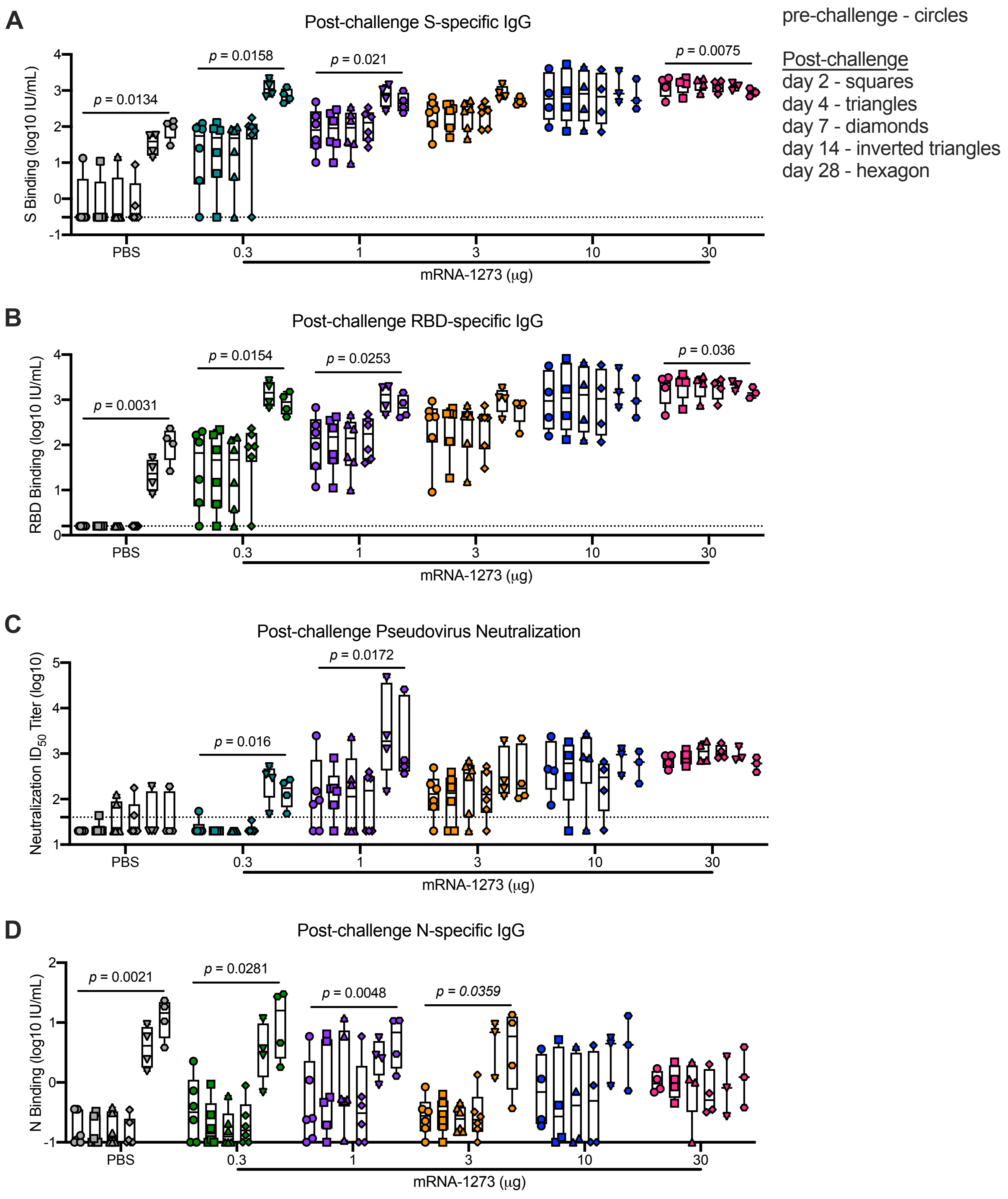


**Fig. S8. Humoral antibody responses following SARS-CoV-2 challenge in mRNA-1273-immunized NHP.** Rhesus macaques were immunized and challenged according to Fig. S1C. Sera collected 4 weeks post-boost, immediately prior to challenge, and days 2, 4, 7, 14, and 28 post-challenge were assessed for SARS-CoV-2 S-specific (A), RBD-specific (B), and N-specific (D) IgG by MULTI-ARRAY ELISA and SARS-CoV-2 D614G lentiviral-based pseudovirus neutralization (C). Boxes and horizontal bars denote the IQR and medians, respectively; whisker end points are equal to the maximum and minimum values. Symbols represent individual NHP. Dotted lines indicate assay limits of detection. Significance between pre-challenge and the final timepoint post-challenge determined by paired t-tests.

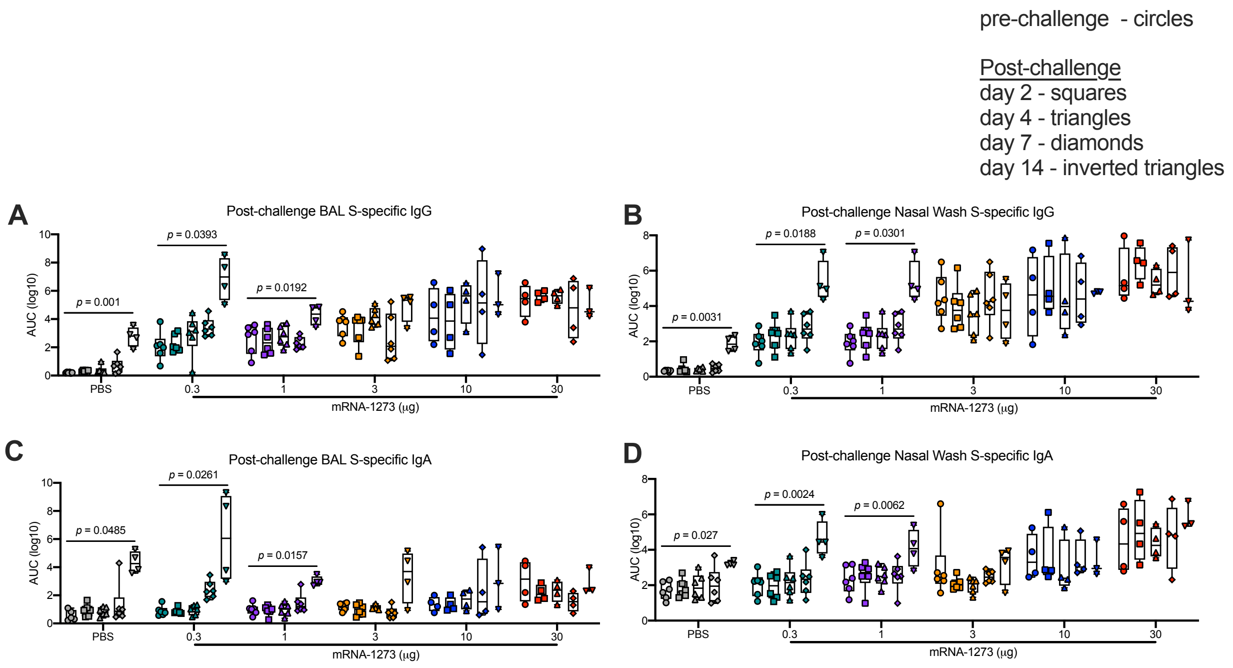


**Fig. S9. Mucosal antibody responses following SARS-CoV-2 challenge in mRNA-1273-immunized NHP.** Rhesus macaques were immunized and challenged according to Fig. S1C. BAL (A, C) and nasal washes (B, D) collected 2 weeks post-boost and days 2, 4, 7, and 14 post-challenge were assessed for SARS-CoV-2 S-specific IgG (A-B) and IgA (C-D) by MULTI-ARRAY ELISA. Boxes and horizontal bars denote the IQR and medians, respectively; whisker end points are equal to the maximum and minimum values. Circles represent individual NHP. Significance between pre-challenge and the final timepoint post-challenge determined by paired t-tests.

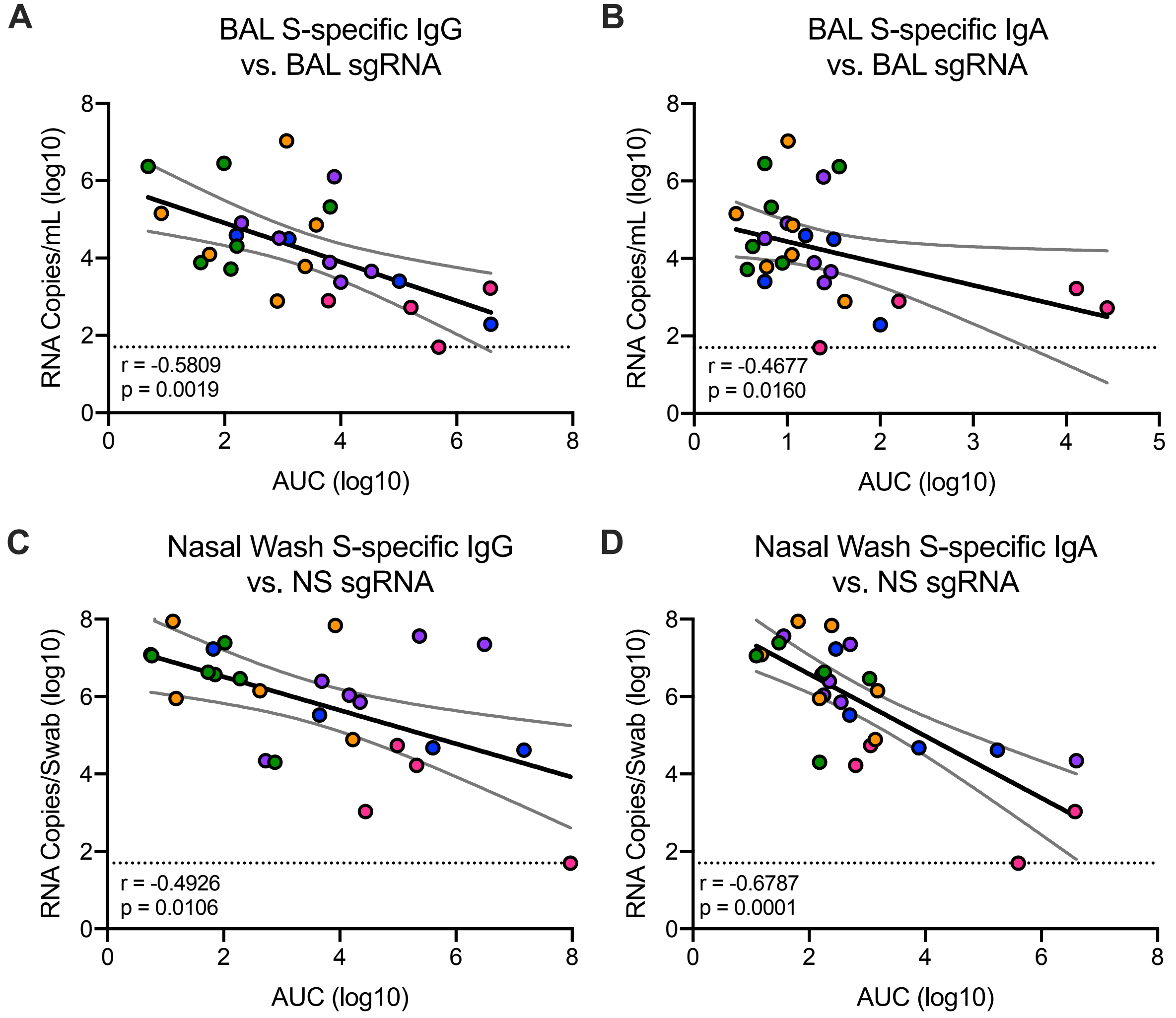


<u>**Fig. S10**</u>**. Correlations of mucosal antibody responses and viral replication.** Rhesus macaques were immunized and challenged according to Fig. S1C. Plots show correlations between BAL (A-B) and NS (C-D) S-specific IgG (A, C) and IgA (B, D) at week 2 post-boost with SARS-CoV-2 N-specific sgRNA in BAL (A-B) and NS (C-D) at day 2 post-challenge. Circles represent individual NHP, where colors indicate mRNA-1273 dose. Dotted lines indicate PCR limit of detection. Black and gray lines indicate linear regression and 95% confidence interval, respectively. 'r' represents Spearman's correlation coefficients, and 'p' the corresponding p-values.

| NHP ID | Post-challenge (Day) | mRNA-1273 (µg) | Inflammation (H&E)[1] | SARS-CoV-2 Ag[2] (Lc[3]; Rmid[4]; Rc[5]) |
|---|---|---|---|---|
| 15D043 | 7 | PBS | ++ | -;+ ;+/- |
| A15V047 | 8 | PBS | ++ | +/-; -; +/ - |
| LP73 | 7 | 0.3 | + | -;-;+/- |
| G45R | 8 | 0.3 | ++ | -;-;- |
| A16V036 | 7 | 1 | + | -;-;- |
| A15V075 | 8 | 1 | + | +/-; -; - |
| 16C282 | 7 | 3 | + | -;-;- |
| 15D025 | 8 | 3 | + | -;-;+/- |
| A15V037 | 7 | 10 | +/- | -;-;+/- |
| 16C225 | 7 | 30 | +/- | -;-;- |

**Table S1. Assessment of lung inflammation and viral antigen following SARS-CoV-2 challenge of mRNA-1273-immunized NHP.**

Lung tissue was evaluated for the presence of inflammation and SARS-CoV-2 viral antigen.

[1] Inflammation scoring: +/- = minimal to mild, + = mild to moderate, ++ = moderate to severe

[2] Immunohistochemistry SARS-CoV-2 antigen (Ag): - = no detection of virus Ag, +/- = rare/occasional Ag+ foci, + = multiple Ag+ foci

[3] Lc: left caudal lung lobe

[4] Rmid: right middle lung lobe

[5] Rc: right caudal lung lobe

**A**

| Outcome | Week 4 Post-boost Antibody Measurement | Linear Regression: Univariate, Outcome ~ Antibody measure: Beta Antibody | p-value Antibody | AdjR$^2$ | Linear Regression: Multivariate, Outcome ~ Antibody measure + dose: Beta Antibody | p-value Antibody | Beta dose | p-value dose | Adj R$^2$ | Antibody measurement meets criteria for Potential CoP? |
|---|---|---|---|---|---|---|---|---|---|---|
| BAL Day 2 sgRNA_N | S-specific IgG | -0.885 | 0.001 | 0.35 | -0.581 | 0.093 | -0.554 | 0.214 | 0.36 | Yes |
| | RBD-specific IgG | -0.936 | <0.001 | 0.38 | -0.659 | 0.050 | -0.513 | 0.223 | 0.39 | Yes |
| | ACE2 Binding Inhibition | -0.957 | <0.001 | 0.49 | -1.029 | 0.008 | 0.122 | 0.812 | 0.47 | Yes |
| | Lentivirus Pseudovirus Neutralization | -0.844 | 0.005 | 0.25 | -0.252 | 0.606 | -0.849 | 0.153 | 0.29 | Yes |
| | VSV Pseudovirus Neutralization | -0.758 | 0.031 | 0.19 | -0.256 | 0.553 | -1.066 | 0.103 | 0.27 | Yes |
| | Live Virus Neutralization | -1.218 | 0.001 | 0.41 | -0.933 | 0.053 | -0.515 | 0.371 | 0.41 | Yes |
| NS Day 2 sgRNA_N | S-specific IgG | -0.966 | 0.003 | 0.29 | -0.499 | 0.222 | -0.851 | 0.115 | 0.34 | Yes |
| | RBD-specific IgG | -1.047 | 0.001 | 0.34 | -0.645 | 0.105 | -0.745 | 0.142 | 0.37 | Yes |
| | ACE2 Binding Inhibition | -1.098 | <0.001 | 0.47 | -1.065 | 0.021 | -0.056 | 0.928 | 0.44 | Yes |
| | Lentivirus Pseudovirus Neutralization | -1.157 | 0.001 | 0.36 | -0.770 | 0.168 | -0.555 | 0.399 | 0.35 | Yes |
| | VSV Pseudovirus Neutralization | -1.027 | 0.023 | 0.22 | -0.133 | 0.789 | -1.898 | 0.017 | 0.41 | No |
| | Live Virus Neutralization | -1.497 | 0.003 | 0.37 | -0.718 | 0.208 | -1.41 | 0.056 | 0.46 | marginal |

**B**

| | | Linear Regression | | | | | | | | |
|---|---|---|---|---|---|---|---|---|---|---|
| | | Univariate<br>Outcome ~ T-cell measure | | | Multivariate<br>Outcome ~ T-cell measure + S-specific IgG | | | | | T-cell measurement predictive after adjusting for S-specific IgG? |
| **Outcome** | **Week 2 Post-boost T cell Measurement** | **Beta predictor** | **p-value predictor** | **Adj $R^2$** | **Beta predictor** | **p-value predictor** | **Beta S-IgG** | **p-value S-IgG** | **Adj $R^2$** | |
| BAL Day 2 sgRNA_N | IL-21 | -4.527 | 0.080 | 0.09 | -0.471 | 0.812 | -1.326 | <0.001 | 0.54 | No |
| | CD40L | -1.284 | 0.008 | 0.24 | -0.313 | 0.459 | -1.226 | <0.001 | 0.55 | No |
| | Any Th1 | -17.106 | 0.004 | 0.28 | -4.874 | 0.364 | -1.184 | 0.001 | 0.55 | No |
| | Any Th2 | -74.749 | 0.011 | 0.22 | -12.775 | 0.622 | -1.265 | <0.001 | 0.54 | No |
| NS Day 2 sgRNA_N | IL-21 | -10.403 | 0.000 | 0.46 | -7.801 | 0.002 | -0.851 | 0.011 | 0.59 | Yes |
| | CD40L | -2.151 | 0.000 | 0.53 | -1.638 | 0.002 | -0.647 | 0.071 | 0.57 | Yes |
| | Any Th1 | -22.922 | 0.001 | 0.38 | -14.489 | 0.045 | -0.816 | 0.051 | 0.45 | marginal |
| | Any Th2 | -67.906 | 0.055 | 0.11 | -4.649 | 0.898 | -1.291 | 0.007 | 0.34 | No |

**Table S2. Summary of linear regression models examining relationships.** (A) Antibody measurements at week 4 post-boost and sgRNA in BAL or NS, with or without adjustment for dose. (B) T cell subsets at week 2 post-boost and sgRNA in BAL or NS, with or without adjustment for S-specific IgG. In all models, sgRNA, antibody measures, and dose are modeled on the $log_{10}$ scale. Antibody measures are considered to meet the criteria for potential correlates of protection from high sgRNA if they are significantly associated with sgRNA univariately and if dose is not statistically significant after adjustment for that potential antibody measure in a multivariate regression model. Gray shading for S-specific IgG represents the pre-specified primary correlate of interest.
